# A nitrogen source-regulated microprotein confers an alternative mechanism of G1/S transcriptional activation in budding yeast

**DOI:** 10.1101/2020.04.20.033787

**Authors:** Sylvain Tollis, Jaspal Singh, Yogitha Thattikota, Roger Palou, Ghada Ghazal, Jasmin Coulombe-Huntington, Xiaojing Tang, Susan Moore, Deborah Blake, Eric Bonneil, Catherine A. Royer, Pierre Thibault, Mike Tyers

**Affiliations:** Institute of Biomedicine, University of Eastern Finland, Kuopio, Finland; Institute for Research in Immunology and Cancer, University of Montreal, Montreal, Quebec, Canada; Lunenfeld-Tanenbaum Research Institute, Mount Sinai Hospital, Toronto, Ontario, Canada; Department of Biological Sciences, Rensselaer Polytechnic Institute, Troy, NY, USA

**Keywords:** Saccharomyces cerevisiae, genetic screen, G1/S transcription, Start, transcription factor, nutrient limitation, microprotein, evolution, quantitative microscopy

## Abstract

Commitment to cell division at the end of G1 phase, termed Start in the budding yeast *Saccharomyces cerevisiae,* is strongly influenced by nutrient availability. To identify new dominant activators of Start that might operate under different nutrient conditions, we screened a genome-wide ORF overexpression library for genes that bypass a Start arrest caused by absence of the G1 cyclin Cln3 and the transcriptional activator Bck2. We recovered a hypothetical gene *YLR053c,* renamed *NRS1* for Nitrogen-Responsive Start regulator 1, which encodes a poorly characterized 108 amino acid microprotein. Endogenous Nrs1 was nuclear-localized, restricted to poor nitrogen conditions, induced upon mTORCl inhibition, and cell cycle-regulated with a peak at Start. *NRS1* interacted genetically with *SWI4* and *SWI6,* which encode subunits of the main G1/S transcription factor complex SBF. Correspondingly, Nrs1 physically interacted with Swi4 and Swi6 and was localized to G1/S promoter DNA. Nrs1 exhibited inherent transactivation activity and fusion of Nrs1 to the SBF inhibitor Whi5 was sufficient to suppress other Start defects. Nrs1 appears to be a recently evolved microprotein that rewires the G1/S transcriptional machinery under poor nitrogeny conditions.

**Author Summar:** Unicellular microorganisms must adapt to ever-changing nutrient conditions and hence must adjust cell growth and proliferation to maximize fitness. In the budding yeast *Saccharomyces cerevisiae*, commitment to cell division, termed Start, is heavily influenced by nutrient availability. Our understanding of how Start is activated is based mainly on experiments carried out under rich nutrient conditions. To identify potential new Start regulators specific to poor nutrient environments, we screened for genes able to bypass a genetic Start arrest caused by loss of the G1 cyclin Cln3 and the transcriptional activator Bck2. This screen uncovered *YLR053c*, which we renamed *NRS1* for Nitrogen-Responsive Start regulator. Sequence analysis across yeast species indicated that Nrs1 is a recently-evolved microprotein. We showed that *NRS1* is nutrient- and cell cycle-regulated, and directly binds the main G1/S transcription factor complex SBF. We demonstrated that Nrs1 has an intrinsic trans-activation activity and provided genetic evidence to suggest that Nrs1 can bypass the requirement for normal Cln3-dependent activation of G1/S transcription. These results uncover a new mechanism of Start activation and illustrate how microproteins can rapidly emerge to rewire fundamental cellular processes.

## Introduction

All organisms have evolved adaptive regulatory mechanisms to optimize fitness in the face of ever-changing environmental conditions. This ability to adapt is particularly important for unicellular organisms, which lack the capacity to establish the internal homeostatic environments of metazoan species. In the budding yeast *Saccharomyces cerevisiae*, different carbon and nitrogen sources can dramatically affect the rates of cell growth and division, as well as developmental programs [1]. Yeast cells commit to division at the end of G1 phase, an event referred to as Start [2, 3]. In order to pass Start, cells must achieve a characteristic critical size threshold that dynamically adjusts to changing nutrient availability thereby optimizing competitive fitness [3]. How nutrient conditions modulate the growth and division machinery at the molecular level is still largely unknown.

Start initiates a complex G1/S transcriptional program of ~200 genes that encode proteins necessary for bud emergence, DNA replication, spindle pole body duplication and other processes. This program is controlled by two transcription factor complexes, SBF (Swi4/6 Cell Cycle Box or SCB binding factor) and MBF (*M/ui* Cell Cycle Box or MCB binding factor), each comprised of related DNA-binding proteins, Swi4 and Mbpl respectively, coupled to a common regulatory subunit Swi6 [4, 5]. Individually Swi4 and Mbpl are not essential but a double *swi4Δ mbp1Δ* mutant is inviable [6], consistent with the significant overlap between SBF and MBF binding sites in G1/S promoters [7–10]. In pre-Start cells that have not achieved the critical cell size, SBF is inhibited by the Whi5 transcriptional repressor [11–13]. At Start, the G1 cyclin (Cln)-Cdc28 protein kinases phosphorylate both SBF and Whi5 to disrupt the SBF-Whi5 interaction and trigger Whi5 nuclear export [12–14]. The upstream G1 cyclin Cln3 is thought to initiate a positive feedback loop in which SBF-dependent expression of *CLN1/2* further amplifies Cln-Cdc28 activity and thus SBF activation [15, 16]. The expression of Cln3 itself does not rely on SBF-dependent positive feedback [17]. Although *CLN3* was isolated as a potent dose-dependent activator of Start [18, 19], the size-dependent mechanism whereby Cln3-Cdc28 initiates the SBF positive feedback loop remains uncertain [20, 21]. *CLN3* is essential only in the absence of other parallel activators of Start, most notably *BCK2,* which encodes a general transcriptional activator [22, 23].

Connections between the main yeast nutrient signalling conduits, the cell size threshold and Start have been established. Activation of the protein kinase A (PKA) pathway by glucose represses *CLN1* and other G1/S transcripts, which may increase the cell size threshold in rich nutrients [24, 25]. The TOR signalling network, which controls many aspects of cell growth including the rate of ribosome biogenesis, has been linked to the size threshold [11, 26–28]. The TORCl complex phosphorylates and activates the effector kinase Sch9 and the master transcription factor Sfpl to activate ribosomal protein (RP) and ribosome biogenesis (Ribi) genes [26, 27, 29]. Deletion of either *SFP1* or *SCH9* abrogates the carbon source-dependent control of cell size [26]. The RimlS kinase, which is active under respiratory growth conditions in poor carbon sources, suppresses the Cdc55 phosphatase that dephosphorylates Whi5, and thereby contributes to Whi5 inactivation even when Cln3 activity is low [30]. Poor nutrient conditions also increase the expression of the G1/S transcription factors and thereby activate Start at a smaller cell size [31]. Notably though, a *cln3Δ bck2Δ whi5Δ* triple mutant is completely viable and still responds to nutrient cues, suggesting that nutrient regulation of Start may be partly independent of the Cln3-Bck2-Whi5-SBF axis [12, 26].

To identify activators of Start that might act in parallel to the central Cln3-Bck2-Whi5 pathway, we screened for dosage suppressors of the lethal *cln3Ll bck2Ll* Start arrest phenotype [12, 13, 32, 33]. The screen identified *YLR053c,* a poorly characterized hypothetical gene that encodes a recently evolved 108 amino acid microprotein, which we renamed *NRS1* for <u>N</u>itrogen-<u>R</u>esponsive <u>S</u>tart regulator 1. Nrs1 protein was only observed in poor nitrogen conditions and was cell cycle-regulated with peak nuclear localization at Start. Nrs1 interacted genetically and physically with SBF and caused a small size phenotype when overexpressed in wild type cells. Nrs1 exhibited intrinsic trans-activation activity and direct fusion of Nrs1 to Whi5 was sufficient to reduce cell size in rich carbon sources and rescue the *cln3Δ bck2Δ* lethality. These results demonstrate that the recently evolved Nrs1 microprotein allows cells to adapt to poor nitrogen conditions by rewiring the Start transcriptional machinery

## Result

### A genome-wide screen identifies overexpression of YLR053c as a rescue of cln3Δ bck2Δ lethality

We constructed a *cln3Δ bck2Δ. Whi5::GAL1-WHI5* strain that is viable in glucose medium but not in galactose due to conditional *WHI5* expression [12, 13]. We used SGA methodology to cross this query strain to an array of 5280 strains that each of contained a *GAL1-GST-ORF* 2 μm high-copy plasmid [34–36], and assessed the growth of the resulting array on galactose medium (Figure 1A, B). Three replicates of the screen identified 12 genes that reproducibly rescued the *cln3Δ bck2Δ* lethal Start arrest (Supplemental Table 1). These genes included the G1 cyclins *CLN1* and *CLN3* but not other known Start activators such as *CLN2, BCK2, 5WI4* and *5WI6*, possibly due to the high level of *WHI5* expression used in our screen, and/or overexpression toxicity of particular genes. We recovered 10 genes not known to be Start regulators. Notably, the short hypothetical ORF *YLR053c* restored *cln3Δ bck2Δ* growth to the same extent as *CLN3,* as validated by direct transformation of the query strain with *GAL1-YLR053c* and *GAL1-CLN3* constructs (Figure 1e). Other prospective high copy rescue genes, such as YEA4 (Figure 1e), could not be confirmed by direct transformation.

**Figure 1.**
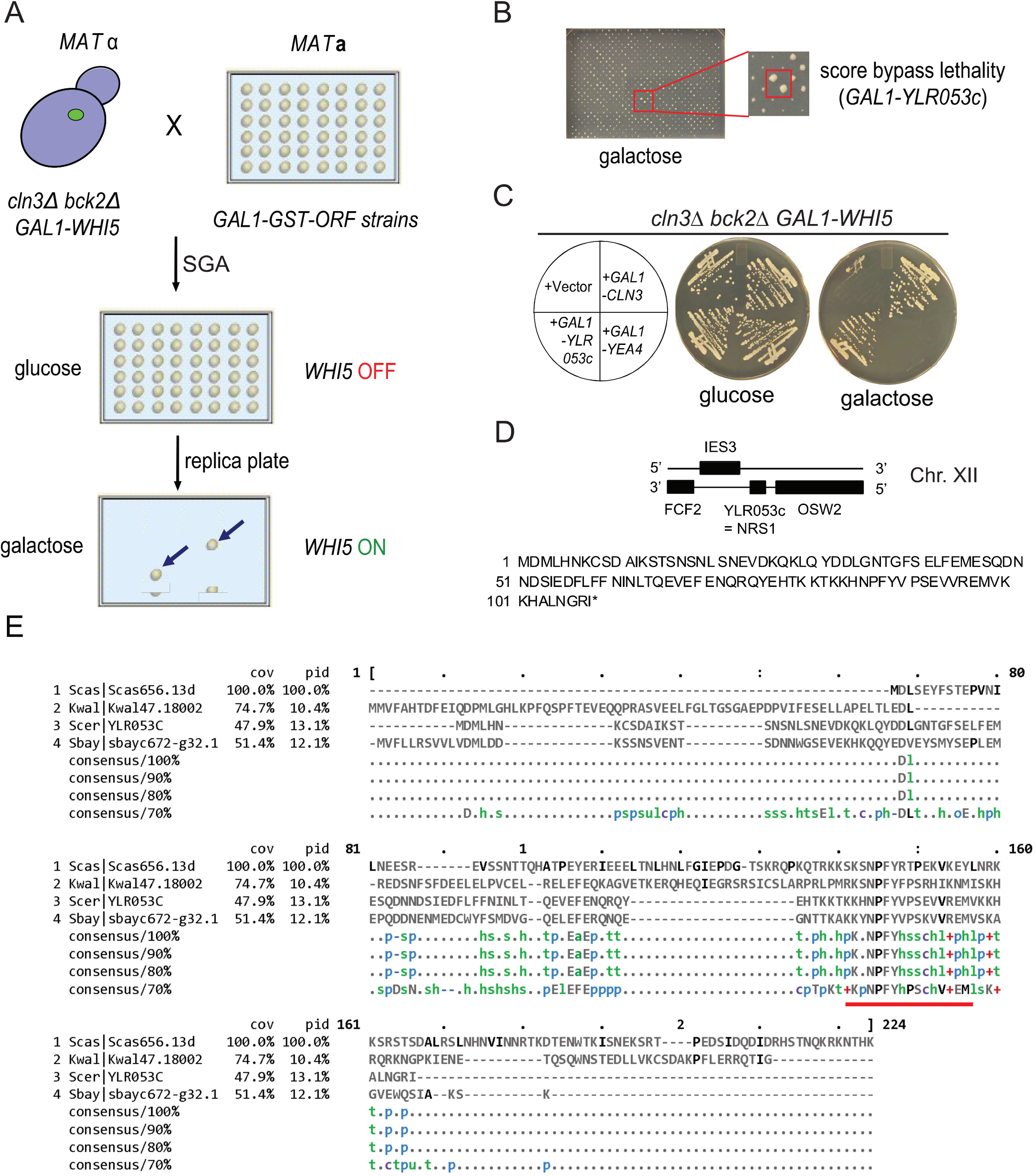
A genome-wide screen identifies *YLR053c/NRS1* as a dosage suppressor of *cln3Δ bck2Δ* lethality. **(A)** Schematic of SGA genetic screen for dosage suppressors of *cln3Δ bck2Δ* lethality. **(B)** Representative screen plate scored for growth on galactose medium. Red box, *GAL1-YLR053c.* **(C)** Comparison of growth at 30°C for the indicated strains streaked onto either glucose or galactose medium. *YEA4* was a candidate hit that did not validate. **(D)** Chromosomal region around *YLR053c/NRS1* on Chr. XII and translated 108 amino acid (12.7 kDa) protein sequence. **(E)** Ylr053c/Nrs1 protein sequence in S. *cerevisiae* (top) aligned with sequences of other yeast species. Conservation of a KKXNPFYVPSXVVREMV motif at the C-terminus is indicated by a red bar.

*YLR053c* encodes a 108 residue microprotein that is only poorly characterized. Microproteins are often encoded by newly evolved proto-genes that are thought to form a genetic reservoir that fuels adaptive evolution [37]. To gain insight into the *YLR053c* locus evolution, we aligned the Ylr053c protein sequence from *S. cerevisiae* with predicted orthologs from other yeast species (Figure 1D, E). To estimate the extent to which *YLR053c* locus evolved across these species, we used the standard dN/dS metric that measures the ratio of single DNA site substitution rates at nonsynonymous codons (dN) versus synonymous codons (dS). The *YLR053c* locus appears to have evolved relatively rapidly within the *5accharomyces* sensu stricto group, with a dN/dS ratio in the 98^th^, 79^th^ and 90^th^ percentile in *S. mikatae, S. bayanus* and *S. castellii,* respectively as compared with *S. cerevisiae* (Figure S1A). In comparison, the evolution of other core Start regulators was closer to the genome median, with dN/dS ratios in the 74^th^, 77^th^ and 58^th^ percentile for *WHI5,* 69^th^, 75^th^ and 31^st^ percentile for *SWl4,* and 82^nd^, 51^st^ and 43^rd^ percentile for *5Wl6*. A 17 amino acid sequence at the Ylr053c C-terminal region was conserved even in more distant yeasts sue as *K. waltii* (Figure 1E). From this point forward, we refer to *YLR053c* as *NRS1* for reasons described below.

### Nrs1 is induced by rapamycin and nitrogen limitation

To understand the function of Nrs1 we first sought to characterize its endogenous expression. We performed scanning Number and Brightness (sN&B) confocal microscopy to localize and quantify an Nrs1-GFPmut3 fusion protein at the subcellular scale in live wild type cells under a range of conditions [31, 38]. GFPmut3, a monomeric fast-folding GFP mutant [39], will be referred to here as GFP for brevity. Nrs1-GFP was not detected in cells grown on either SC + 2% glucose or SC + 2% raffinose medium (Figure 2A), consistent with previous analysis of *YLR053/NRS1* mRNA levels [40]. In contrast, Nrs1-GFP was readily detected in the nucleus of cells grown overnight to mid log-phase in nitrogen-limited (YNB + 0.4% praline + 2% glucose, abbreviated YNB+Pro) medium (Figure 2A). This result was consistent with published genome-wide transcriptome analyses under various nutrient conditions [41]. Induction of Nrs1 upon the switch to nitrogen-limited YNB+Pro medium required a long incubation time, as Nrs1-GFP signal above autofluorescence background could not be detected after 7 h in YNB+Pro but was clearly visible after 22 h growth to mid-log phase in YNB+Pro (Figure S1B). Inhibition of TORCl with 100 nM rapamycin induced Nrs1-GFP expression and nuclear localization within 1 h in rich SC + 2% glucose medium (Figure 2A; Figure S1C}. These expression patterns were confirmed with an independent fusion of Nrs1 to wild type GFP {Figure S1D). Hyperosmotic stress, oxidative stress and DNA damage did not induce Nrs1 expression in SC+ 2% glucose medium {Figure S1E, SlF). lmmunoblot analysis of strains bearing a MYC epitope-tagged version of Nrs1 in different carbon and nitrogen sources, as well as in rapamycin-treated and saturated cultures, confirmed the specific expression of Nrs1 only under conditions of nitrogen limitation or TORCl inhibition (Figure 2B). Consistently, the *NRS1* promoter contains binding sites for the Gln3 transcription factor, which activates gene expression in response to nitrogen limitation and rapamycin treatment [42, 43]. In accord with its genetic role at Start and expression pattern, we renamed the *YLR053c* gene *NRS1* for <u>N</u>itrogen-<u>R</u>esponsive <u>S</u>tart regulator.

**Figure 2.**
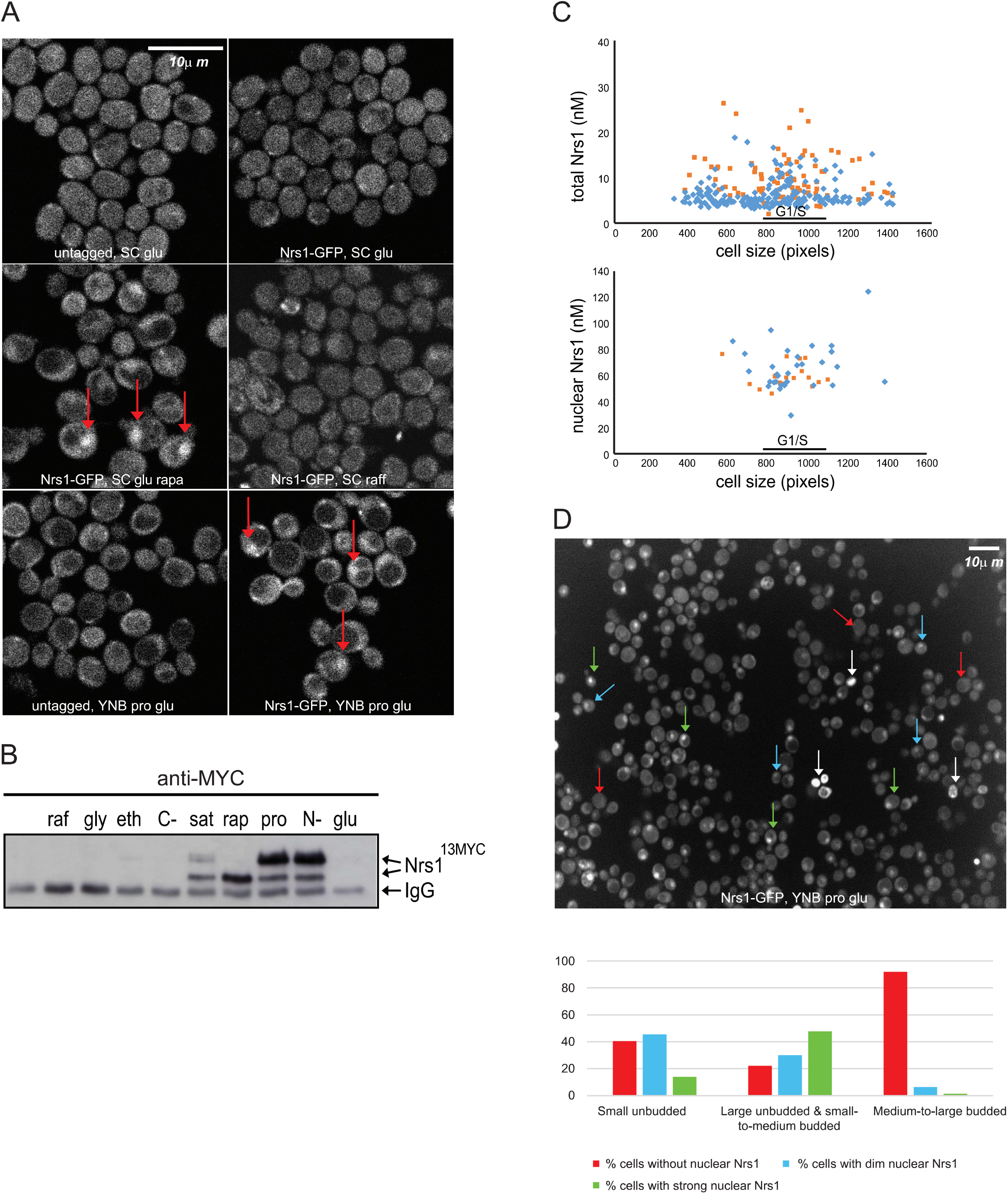
Nitrogen limitation and rapamycin treatment induce cell cycle-regulated Nrs1 expression. **(A) sN&B** images of untagged and *NRS1-GFP* strains. Cells were grown and imaged in SC+ 2% glucose with or without 100 nM rapamycin, SC+ 2% raffinose or nitrogen-limited medium (YNB +0.4% praline+ 2% glucose+ His, Leu, Met, Ura; labelled YNB+Pro glu), as indicated. Scale bar is 10 μm. The same intensity scale was used for all conditions. **(B)** Abundance of Nrs1^13MYC^ in various nutrient conditions as determined by immunoblot of anti-MYC immune complexes. raf, 2% raffinose; gly, 2% glycerol; eth, 2% ethanol; C-, no carbon source; sat, saturated culture; rap, 100 nM rapamycin; pro, 0.4% praline; N-, no nitrogen source; glu, 2% glucose. lgG indicates antibody light chain. **(C)** Absolute Nrs1 concentration in single cells grown and imaged in nitrogen-limited medium (YNB+Pro glu) as a function of cell size, as determined by sN&B. Blue and orange dots represent individual cells from two different experiments. The typical size range of cells at the G1/S transition (800-1000 pixels, corresponding to 27-38 fl) is indicated. Cell-averaged total Nrs1 concentration (top) and nuclear concentration where Nrs1 nuclear localization was evident (bottom) are shown. Infrequent small cells with high Nrs1 levels had high autofluorescence and no nuclear localization of the signal. **(D)**Example of high-content confocal image of Nrs1-GFP cells grown to log phase in nitrogen-limited medium. Arrows indicate example cells with strong (green), dim (blue) or no nuclear Nrs1 (red) or inviable cells (white). The histogram summarizes Nrs1 signals in small unbudded (708 cells), large unbudded & small-t medium budded (599 cells) and medium-to-large budded cells (186 cells) from 18 confocal images. Inviable cells with high autofluorescence, out-of-focus cells for which the budding pattern could not be ascertained and regions in which illumination was not homogeneous were not scored.

### Nrs1 abundance peaks at the G1/S transition

We quantified Nrs1-GFP levels in asynchronous populations of live cells grown in nitrogen-limited YNB medium with our custom sN&B analysis software [31]. Single cell-averaged Nrs1-GFP levels were between 5 and 20 nM. Higher levels were observed in cells close to the typical critical size at the end of G1 phase, where the predominant nuclear Nrs1-GFP signal corresponded to 60-80 nM (Figure 2C). These Nrs1 nuclear levels at Start were comparable to Swi4 (50-100 nM), Whi5 (100-120 nM) and Swi6 (130-170 nM) levels determined previously by the same method [31]. We further confirmed Nrs1-GFP expression and nuclear localization principally in large unbudded and small-to-medium budded cells in high-content images acquired by standard confocal microscopy (Figure 2D). The cell cycle-regulated expression pattern of Nrs1 protein suggested that it plays a role at Start, consistent with the suppression of the *cln3Δ bck2Δ* arrest by *NRS1* overexpression.

### NRS1 genetically interacts with SBF and other Start regulators

To investigate the molecular mechanism by which Nrs1 promotes Start, we examined genetic interactions of *NRS1* with the main known Start regulators. Overexpression of *NRS1* from the *GAΔ* promoter, either integrated at the *NRS1* locus or from a 2 μm high copy plasmid [44], caused a pronounced small size phenotype, in agreement with its putative role as a Start activator (Figure 3A, B). To interrogate the genetic requirements for this size phenotype, we transformed deletion mutants of known Start regulators with the *GAL1-NRS1* high copy plasmid and examined cell size epistasis. *NRS1* overexpression almost entirely rescued the large size phenotype of a *cln3Δ* mutant and partially rescued the large size of a *bck2Δ* mutant (Figure 3B). A *whi5Δ* mutant was epistatic to *NRS1* overexpression, whereas a *nrs1Δ* deletion did not exacerbate the larger size caused by *WHI5* overexpression and only modestly increased the size of a *whi5Δ* mutant (Figure 3C). Notably, the small size caused by *NRS1* overexpression was abrogated in *swi6Δ* and *swi4Δ* mutant strains (Figure 3D). This requirement for full SBF function was further demonstrated with a temperature sensitive *swi4* strain in which Swi4 binding to Swi6 is altered [45], grown at the semi-permissive temperature of 30°C (Figure 3E).

**Figure 3.**
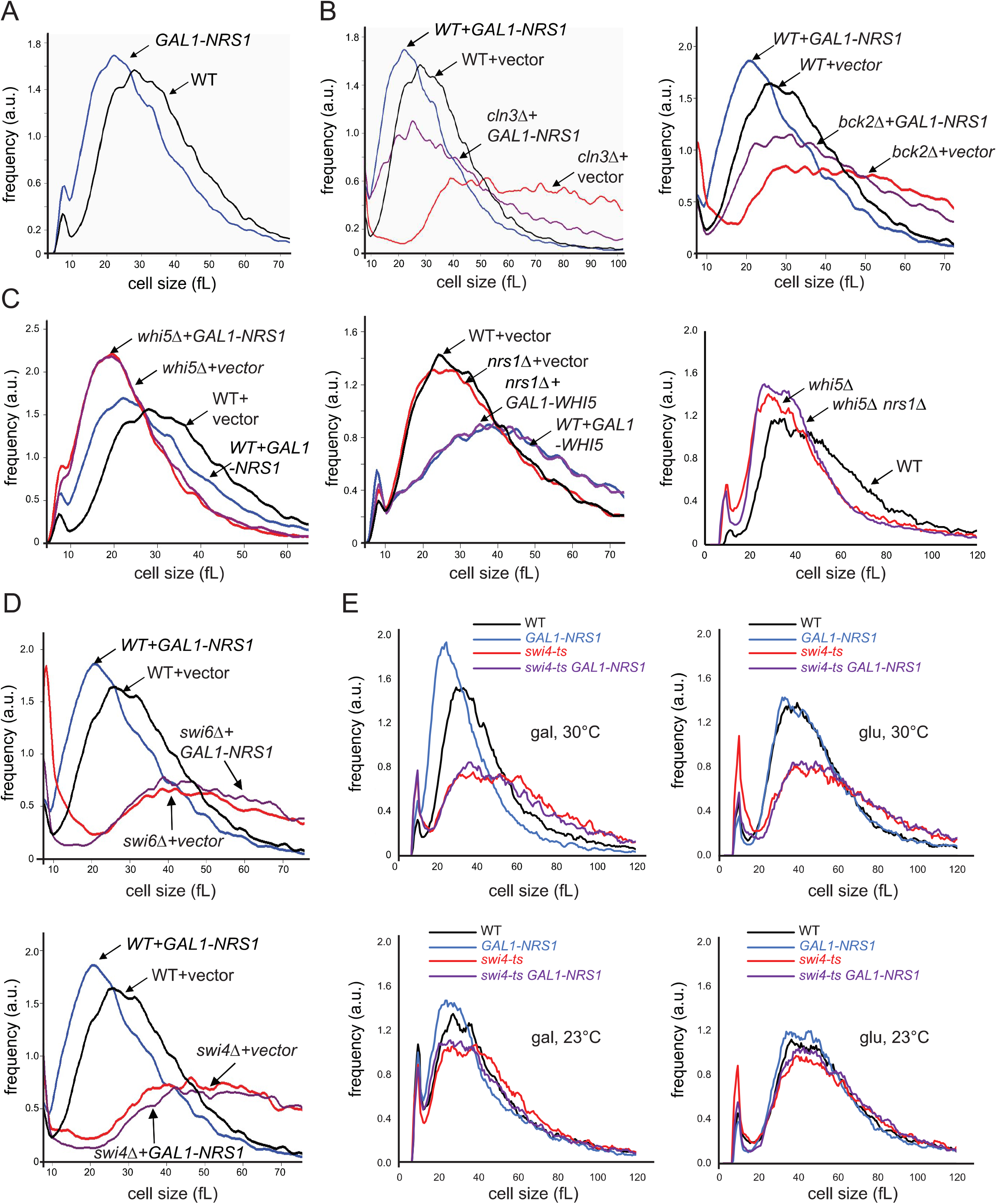
*NRS1* is genetically upstream of SBF. Cell size distributions were determined for the indicated genetic combinations. (A) *NRS1* overexpression alone. (B) *NRS1* overexpression with either *cln3Δ* and *bck2Δ* mutations. (C) *NRS1* overexpression and *nrs1Δ* mutation with either *WHI5* overexpression or *whi5Δ* mut ation. (D) *NRS1* overexpression with either *swi4Δ* or *swi6Δ* mutations. (E) *NRS1* overexpression with a *swi4-ts* mutation. Strains were transformed with either *GAL1-NRS1, GAL1-WHI5* or empty vector control high copy plasmids as indicated. Strains bearing galactose-regulated constructs and associated controls were induced for 6 h in SC+ 2% galactose before size determination. Wild type plots shown in panels B, C and D (left, middle) represent the same measurement.

In contrast to its overexpression, deletion of *NRS1* did not detectably affect overall growth rate or cell size in the S288C (BY4741) laboratory strain background, regardless of whether cells were grown in either nitrogen-replete or nitrogen-limited medium (Figure S2A-D). Moreover, in competitive growth experiments, *nrs1Δ* and wild type cells had indistinguishable fitness (Figure S2E). Because the small size phenotype caused by *NRS1* overexpression depended on *SWl4* and *SWl6,* we hypothesized that MBF might partially compensate for loss of *NRS1* function. We therefore generated an *mbp1Δ nrs1Δ* double mutant strain and evaluated growth in nitrogen-rich SC and nitrogen-poor YNB+Pro media. While the single mutant strains had no growth defect in either condition, the *mbp1Δ nrs1Δ* double mutant had a pronounced growth defect that was specific to nitrogen-poor conditions (Figure S2F). We furthermore considered the possibility that laboratory strains may have lost some requirements for Nrs1 in the course of decades-long propagation in artificially rich nutrient conditions. To test this idea, we deleted *NRS1* in the prototrophic wild yeast *5accharomyces boulardii,* which shares >99% of its genome with *S. cerevisiae* including *NRS1* [46]. An *S. boulardii* strain lacking *NRS1* grew slower than wild type cells in nitrogen-poor medium specifically (i.e., YNB+Pro+2% glucose medium not supplemented with amino acids, Figure S2G). Hence, under conditions of nitrogen limitation, *NRS1* promotes growth in genetically crippled contexts in laboratory strains and is required for optimal growth of a wild variant of *S. cerevisiae.* Taken together, these genetic interactions suggested that Nrs1 promotes the G1/S transition by acting upstream of the Whi5-inhibited form of SBF in a manner that is independent of Cln3 and Bck2.

### Endogenous Nrs1 interacts with SBF in vivo and in vitro

We next sought to identify Nrs1 protein interactors under conditions in which Nrs1 was endogenously expressed at physiological levels. We performed immunoaffinity purification on lysates from an endogenously tagged *NRS1*^*13MYC*^ strain and a control untagged strain grown in nitrogen-limited YNB+Pro medium. Analysis of the samples by mass spectrometry identified multiple peptides (Supplemental Table 4) from which the corresponding proteins were identified (> 2 unique peptides, false discovery rate< 1%, Supplemental Table 5). Proteins specific to the Nrs1^13MYC^ sample were identified by subtracting proteins present in the untagged control and filtering hits against the CRAPome database of non-specific interactions [47]. This workflow yielded 7 proteins that were specific to the Nrs1^13MYC^ sample {Supplemental Table 3, Figure S3A). Of these, Nrs1 was represented by 7 peptides {48% coverage), Swi4 by 5 peptides, and Swi6 by 5 peptides; the remaining 4 candidates were represented by only 2 or 3 peptides and thus of lower confidence. This unbiased analysis demonstrated that endogenous Nsrl expressed under physiological conditions interacted with SBF. To confirm this result, we performed co-immunoprecipitation experiments on extracts of rapamycin-treated cells expressing MYC-tagged *NRS1* or *WHI5* alleles from their endogenous promoters or from an untagged control strain (Figure 4A). Nrs1^13MYC^, Whi5^13MYC^, Swi4 and Swi6 were each immunoprecipitated with either anti-MYC or polyclonal antibodies as indicated and then blotted with antibodies against each protein. The specific association of endogenously expressed Nrs1^MYC^ with both Swi4 and Swi6 resembled that of Whi5^MYC^, again suggesting that Nrs1 directly interacted with SBF under physiological conditions.

**Figure 4.**
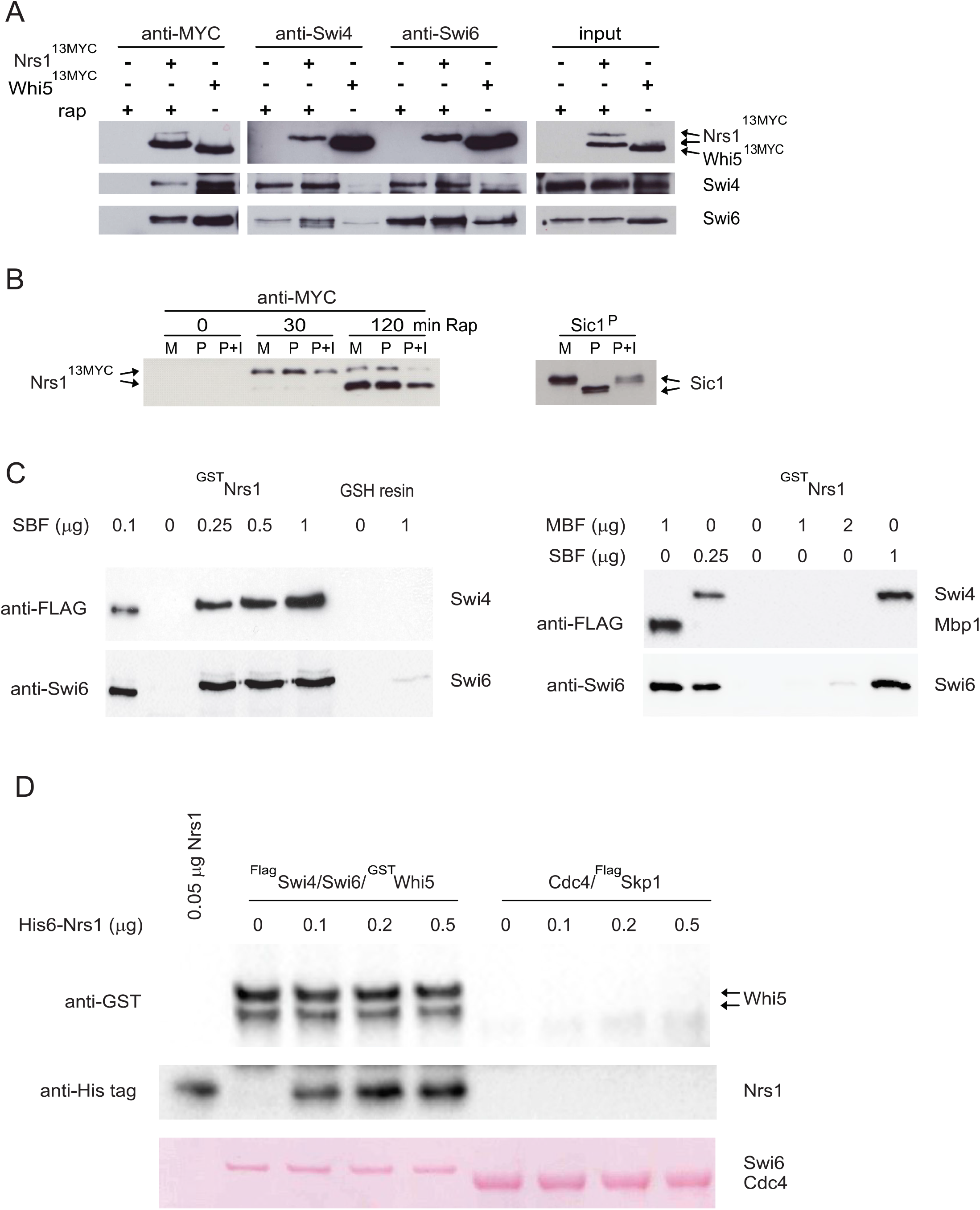
Nrs1 binds to SBF *in vivo* and *in vitro.* **(A)** Nrs1^13MYC^, Swi4, or Swi6 complexes were immunoprecipitated from a rapamycin-treated *NRS1*^*13MYC*^ strain and interacting proteins assessed by immunoblot with the indicated antibodies. Co-immunoprecipitation of Whi5^13MYC^ With Swi4 and Swi6 served as a positive control. **(B)** Nrs1^MYC^ immunoprecipitates from a rapamycin treatment time course were either mock-treated (M, negative control), treated with lambda phosphatase (P) or lambda phosphatase+ phosphatase inhibitors (P+I) prior to detection by anti-MYC immunoblot. *In vitro* phosphorylated recombinant Sicl was used as a control to demonstrate activity of the phosphatase.**(C)** Recombinant ^GST^Nrs1 immobilized on GSH-Sepharose resin was incubated with increasing concentrations of soluble purified SBF (^FLAG^Swi4-Swi6) or MBF (^FLAG^Mbpl-Swi6). Bound proteins were analysed by immunoblot with anti-Swi6 and anti-FLAG antibodies as indicated. GSH-Sepharose resin alone served as a negative control. **(D)** Recombinant ^FLAG^Swi4-Swi6-^GST^whi5 complexes containing 0.1 ug of ^GST^Whi5 were incubated with increasing amounts of recombinant His_6_-tagged Nrs1 (0.1 ug, 0.2ug and 0.Sug, equivalent to 4, 8, and 20 Nrs1:Whi5 molar ratio), immunoprecipitated usig anti-FLAG beads to capture SBF-Whi5 complexes and probed with anti-GST antibody to detect Whi5 and anti-HIS6 antibody to detect Nrs1. Co-immunoprecipitation with an irrelevant recombinant protein complex (Cdc4-^FLAG^Skp1) on anti-FLAG beads served as a negative control for interaction specificity. Ponceau S stain was used to demonstrate equivalent input protein complexes in each lane.

We note that endogenous Nrs1 appeared as a doublet on immunoblots and only the slow-migrating form interacted with SBF (Figure 4A). This slow-migrating form was predominantly induced by nitrogen limitation (Figure 2B) and also appeared first upon rapamycin treatment, before conversion to the fast-migrating form after 2 h (Figure 4B). To investigate the nature of this presumptive post-translational modification **(PTM)** on Nrs1, we treated Nrs1^MYC^ immuno-precipitates with lambda phosphatase but found that this did not affect the slower-migrating species (Figure 4B). We also did not detect Nrs1-derived phosphopeptides in our mass spectrometry profiles (Supplementary Table 4). These results suggest that Nrs1 is regulated by a post-translational control mechanism and that the presumptive modified form preferentially interacts with SBF. While the slow-migrating form of Nrs1 appears not to be caused phosphorylation, we have not been able to determine the nature of this modification.

To determine if the interactions between Nrs1 and SBF were direct, we carried out *in vitro* binding assays with purified recombinant proteins. We titrated recombinant ^FLAG^Swi4-Swi6 and ^FLAG^Mbpl-Swi6 complexes produced in baculovirus-infected insect cells [12] against recombinant ^GST^Nrs1 produced in *E. coli* and immobilized on GSH-Sepharose resin. Swi4 and Swi6 were both capture with ^GST^Nrs1 across the titration series whereas control GSH-Sepharose resin did not bind SBF (Figure 4C). In contrast, ^GST^Nrs1 did not detectably interact with the ^FLAG^Mbpl-Swi6 complex under the conditions tested, suggesting that specificity for Nrs1 is determined by the DNA binding subunit and not the common Swi6 subunit of SBF/MBF. These results demonstrated that endogenous Nrs1 interacts directly and specifically with the Swi4-Swi6 complex, and that no additional factors are needed for this interaction to occur.

### Nrs1 binds to SBF-regulated promoters in vivo

We next assessed the presence of Nrs1^13MYC^ at the SBF-regulated promoters of *CLN2* and *PCLl* by chromatin immunoprecipitation (ChlP). For both genes, we specifically detected SCB-containing promoter sequences by PCR in cross-linked anti-MYC immune complexes purified from cells that expressed Nrs1^13MYC^ from the endogenous *NRS1* locus (Figure 5A). The enrichment of *PCLl* and *CLN2* sequences in Nrs1^13MYC^ immunoprecipitates only in rapamycin-treated cells further demonstrated that Nrs1 specifically binds SBF-regulated G1/S promoter DNA (Figure 5A).

**Figure 5.**
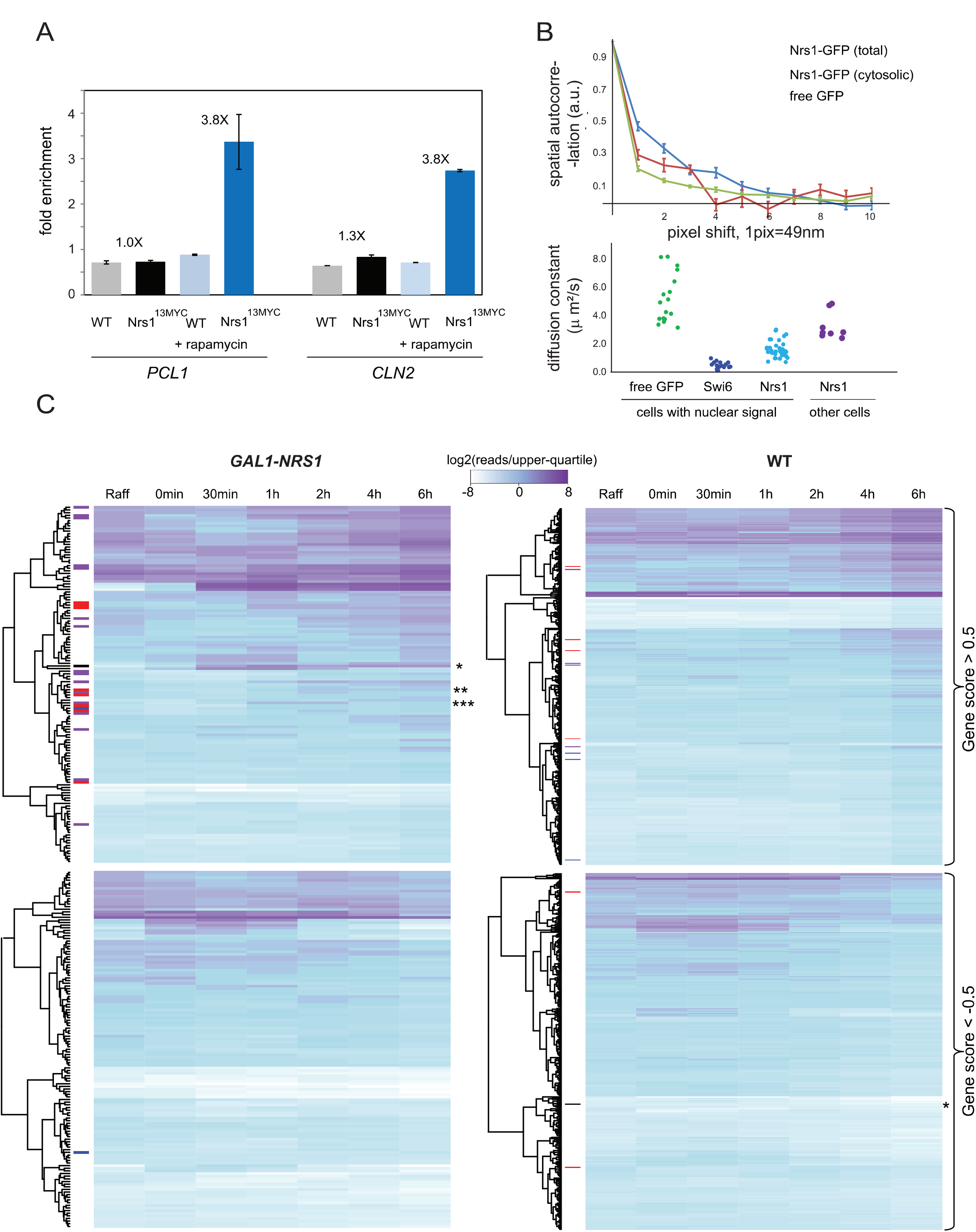
Nrs1 binds to SBF-regulated promoter DNA and activates SBF-driven transcription. **(A)** Anti-MYC chromatin immunoprecipitates from wild type untagged control and *NRS1*^*13MYc*^ strains either untreated or treated with 200 ng/mL rapamycin for 2 h were probed for the presence of *CLN2* and *PCLl* promoter DNA sequences by real time quantitative PCR. Bars indicate the mean fold-enrichment across two replicates, error bars show the standard error on the mean. **(B)** Top panel: RICS vertical correlations from a *NRS1-GFP* strain grown in nitrogen-limited medium (YNB+Pro) as a function of the pixel shift for total and cytosolic Nrs1-GFP pools. RICS correlation for free GFP was used as a control for unconstrained diffusion. Data points show correlation averages over N fields of view (FOV, N=25 for nuclear Nrs1, N=7 for cytosolic Nrs1, N=12 for free GFP). Each FOV contained 1-5 cells. Error bars represent the standard error on the mean. Bottom panel: Scatter plots of fitted diffusion coefficients from individual FOVs of the same strains. Data from Swi6-GFP FOVs served as a control for constrained diffusion. **(C)** Heatmaps of gene expression profiles upon galactose induction of *GAL1-NRS1* and wild type control strains. Genes were selected based on a gene rank score across biological triplicates and values show are average raw log_2_-read counts across triplicates for each timepoint as normalized to the upper quartile of all genes. Genes that were strongly upregulated (gene score> 0.5) and downregulated (gene score < −0.5) are shown. Upregulated and downregulated genes were clustered separately according to the default settings of the R Heatmap function. Positions of NRS1 (*), CLN2 (**) and PCLl (***) are indicated. SBF (red), MBF (blue) and SBF/MBF (purple) target genes are indicated on the left side.

We predicted that Nrs1 binding to chromatin should reduce Nrs1 mobility in the nucleus. To test this hypothesis, we assessed the molecular dynamics of Nrs1-GFP in cells grown in nitrogen-limited medium using Raster Image Correlation Spectroscopy (RICS). The RICS method exploits the hidden time structure of imaging scans, in which adjacent pixels are imaged a few microseconds apart, to quantify the diffusional properties of fluorescent molecules [48–50]. Intensity correlations between pixels shifted along the direction of scanning (horizontal correlations), and along the orthogonal direction (vertical correlations), are averaged over multiple scans. Horizontal and vertical correlations that decay with increasing pixel shift are characteristic of the dynamical properties on 10-100 μs and 5-50 ms timescales, respectively. We found that the vertical RICS correlations of nuclear Nrs1-GFP signal decreased on a slower timescale than either free nuclear GFP or cytosolic Nrs1-GFP signal (Figure 5B). Fitted Nrs1-GFP diffusion coefficients in cells in which Nrs1 was predominantly nuclear were lower compared to cells in which it was mostly cytosolic, and also lower than the diffusion coefficient of free nuclear GFP, a proxy for free diffusion in the nucleus. The Nrs1 diffusion coefficient was close to values for Swi6-GFP which is largely DNA-bound (Figure 5B). This result indicated that Nrs1 associates with a slowly-moving nuclear component, most likely chromatin.

The apparent size epistasis between *whi5Δ* and *GAL1-NRS1* prompted us to ask whether Nrs1 might reduce the association Whi5 with SBF and/or chromatin. However, ChIP analysis of Whi5HA at the *CLN2* and *PCLl* promoters revealed that Whi5HA_promoter interactions were not reduced in the presence of overexpressed *NRS1* (Figure S4A). In addition, *GAL1-NRS1* cells had wild type levels of Whi5 as determined by sN&B quantification (Figure S4B) and *GAL1-NRS1* did not alter Whi5^HA^ association with Swi4 or Swi6 in co-immunoprecipitation experiments (Figure S4C). Nrs1 and Whi5 also did not compete for SBF binding in *in vitro* binding assays with recombinant proteins (Figure S4D). Together, these results obtained using different *in vivo, in vitro,* biochemical and imaging-based approaches suggested that Nrs1 binds SBF at G1/S promoters *in vivo*, but that the binding of Nrs1 does not alter Whi5 interactions with SBF or promoter DNA. These results led us to consider the possibility that Nrs1 might directly activate transcription.

### Ecotopic expression of NRS1 activates the G1/S regulon

To assess the potential role of Nrs1 in transcription, we used RNA-seq to determine the genome-wide transcriptional response to ectopic *NRS1* overexpression. Three biological replicates of *nrs1::GAL1-NRS1* and wild type cells were grown to log-phase in SC+2% raffinose medium, induced with galactose for 6 h, and transcriptional profiles analyzed by RNA-seq. Gene scores were defined based on their expression fold-change before and after 6 h induction, filtering against genes up/down regulated in both strains, hence independent of *NRS1* (see Methods; genes scores provided in Supplementary Table 6). Upregulated genes in the *GAL1-NRS1* samples included *NRS1* itself as expected (rank 1), and most of the 139 SBF/MBF target genes that comprise the G1/S regulon as defined by integration of multiple microarray-based analyses of G1/S regulated genes [10]. These G1/S genes included *CLN2* (rank 12) and *PCLl* (rank 82), the promoters of which were bound by Nrs1 as shown above. Nrs1 also strongly induced the expression of *HO* (rank 10), *RNR1* (rank 15), and *HCMl* (rank 20), which are markers of the G1/S transition. Visualization of the data in a heatmap format illustrated the overall enrichment of SBF/MBF target genes (Figure SC). Amongst the most strongly upregulated genes (score > 0.5), SBF targets were enriched 11-fold compared to random expectation (24 of 89 genes, p=6* 1^−20^) MBF targets were enriched 4-fold (6 of 59 genes, p=0.002) with an overall enrichment of 3.2-fold (30/139 genes, p=2*10^−19^) for the entire G1/S regulon. In the wild type control samples, 10 of the 139 SBF or MBF target genes were strongly upregulated, representing no significant enrichment/depletion compared to random expectations. We note that in addition to recognition of its cognate SCB elements, SBF can also activate genes that only contain MCB elements [51] such that the apparent specificity of Nrs1 for SBF is not undermined by the up regulation of MBF genes in these experiments.

### Nrs1 confers transcriptional activity that can rescue G1/S-transcription deficient mutants

We next asked whether Nrs1 might itself function directly as a transcriptional activator. To test this hypothesis, we constitutively expressed a Nrs1-Gal4 DNA binding domain fusion protein (*GAL4*^*DBD*^ - NRS1) in a reporter strain bearing the *HI53* gene under control of the *GAΔ* promoter. We used *GAL4*^*DBD*^ alone and *GAL4*^*DBD*^ fused to an irrelevant human gene *(UBE2G2)* that potently transactivates as negative and positive controls, respectively. Full length *NRS1,* but not a truncated version that encoded only the conserved C-terminus was able to activate transcription of the *HIS3* reporter and thereby allow cell growth in medium lacking histidine (Figure 6A; Figure S5A).

**Figure 6.**
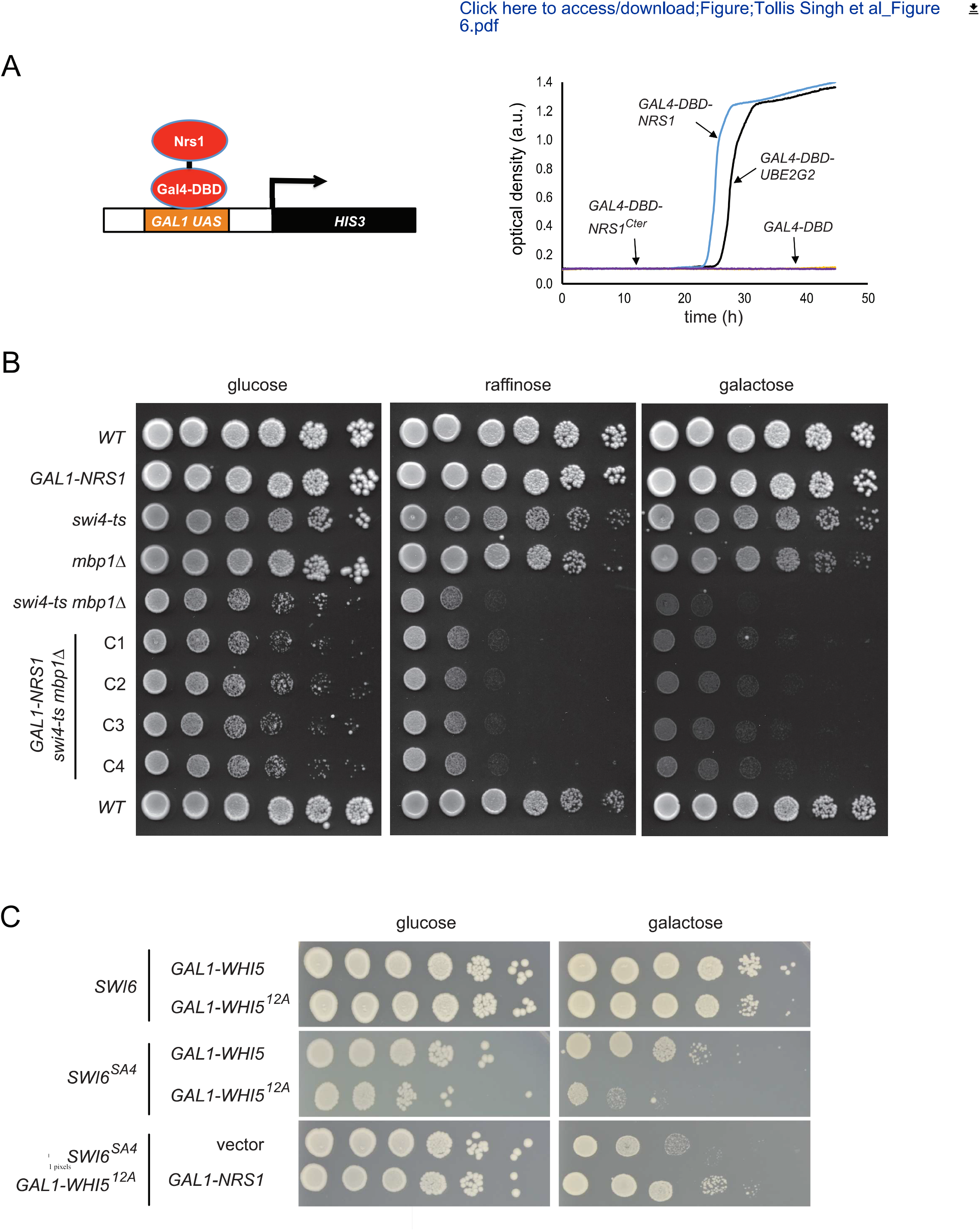
Nrs1 has an inherent transcription activation function that can partially rescue G1/S-transcription deficient mutants. **(A)** Transactivation of a *HI53* reporter by fusion of Nrs1 to the Gal4 DNA binding domain (GAL4^*DBD*^-NRS1) but not by fusion of the Nrs1 C-terminus (*GAL4*^*DBD*^-*NRS1*^*cter*^). *GAL4*^*DBD*^-*UBE2G2 and GAL4*^*DBD*^ constructs served as positive and negative controls, respectively. Growth curves were determined in -His -Trp medium. **(B)** Serial 5-fold dilutions of *NRS1* and *GAL1-NRS1* strains in wild type, *swi4-ts, mbp1Δ and mbp1Δ swi4-ts* backgrounds were spotted onto SC+ 2% glucose, SC+ 2% raffinose and SC+ 2% galactose, and grown for 5 days at 30°C. Cl to C4 indicate four independent clones of the *mbp1Δ swi4-ts GAL1-NRS1* strain. **(C)** Serial 5-fold dilutions of wild type or *swi6Δ* strains transformed with *GALl-WHI5, GAL1-WHI5*^*12A*^, *GAL1-SWI6*^*SA4*^, *GAL1-NRS1* or empty control plasmids or combinations thereof were spotted on rich medium containing 2% glucose or 2% galactose and grown for 2 days at 30°C.

Given the inherent transactivation capacity of Nrs1, we next investigated whether *NRS1* overexpression could suppress the growth defects caused by loss of G1/S activat ors. We first tested a *c/nΔ cln2Δ cln3Δ MET-CLN2* strain, in which G1 cyclin activity is restricted to methionine-free media, with or without a <pGAL1-NRS1> plasmid. Repression of *CLN2* by methionine caused a growth arrest, whether or not *NRS1* was expressed, although ectopic expression of *NRS1* did modestly reduce cell size and slightly increase cell proliferation (Figure S5B). Hence, NRS1 cannot compensate for the total lack of G1-cyclin activity. We then tested whether *NRS1* overexpression could suppress the growth defects of mutants in which G1/S transcription was impaired but in which Cln-Cdc28 activity was preserved. We first tested a *mbpΔ swi4-ts* double mutant [52]. The growth of a *GAL1-NRS1 mbp1Δ swi4-ts* strain was improved with respect to the *mbp1Δ swi4-ts* control at a semi-permissive temperature of 30° in galactose medium (Figure 6B; see Figure S5C for plate images with a 2 day extension of the incubation period), but not in other media or at the fully permissive temperature (Figure S5B). As a further test, we examined NRS1-mediated suppression in strains that expressed *WHI5*^*12A*^ and *SWI6*^*SA4*^ alleles in which all Cdc28 consensus sites are mutated, a combination that abrogates the Cln-Cdc28 mediated relief of SBF inhibition by Whi5 [12, 14). Overexpression of *NRS1* also partially rescued the growth defect of this strain (Figure 6C). These results suggested that Nrs1 provides an alternative mechanism of Start activation by augmenting G1/S transcription in the presence of Whi5-mediated inhibition.

### Nrs1 can bypass Whi5 inhibition of SBF

To test whether physiological levels of Nrs1 targeted to SBF are sufficient to promote Start activation, we constructed a strain expressing a Whi5-Nrs1-GFP fusion protein from the endogenous *WHI5* promoter. Experiments with this strain were carried out in SC complete medium to avoid other possible nitrogen-dependent parallel inputs to Start and focus on Nrs1 function. The chimeric protein was expressed and had the expected molecular mass (Figure S6A) and retained the localization pattern of endogenous Whi5, with about 30-35% of pre-Start cells showing a prominent nuclear signal (Figures 7A, S6B). Cells expressing Whi5-Nrs1-GFP were almost as small as *whi5Δ* cells (Figure 7B). This phenotype was not due to the presence of the GFP tag since size distributions of strains expressing untagged versus GFP-tagged Whi5 were indistinguishable (Figure S6C). Moreover, cells expressing the chimeric protein grew at the same rate as wild type and *whi5Δ* strains (Figure S6D). Strikingly, cells expressing Whi5-Nrs1-GFP were as small as cells that expressed *GAL1-NRS1* in galactose medium (Figure 7C).

**Figure 7.**
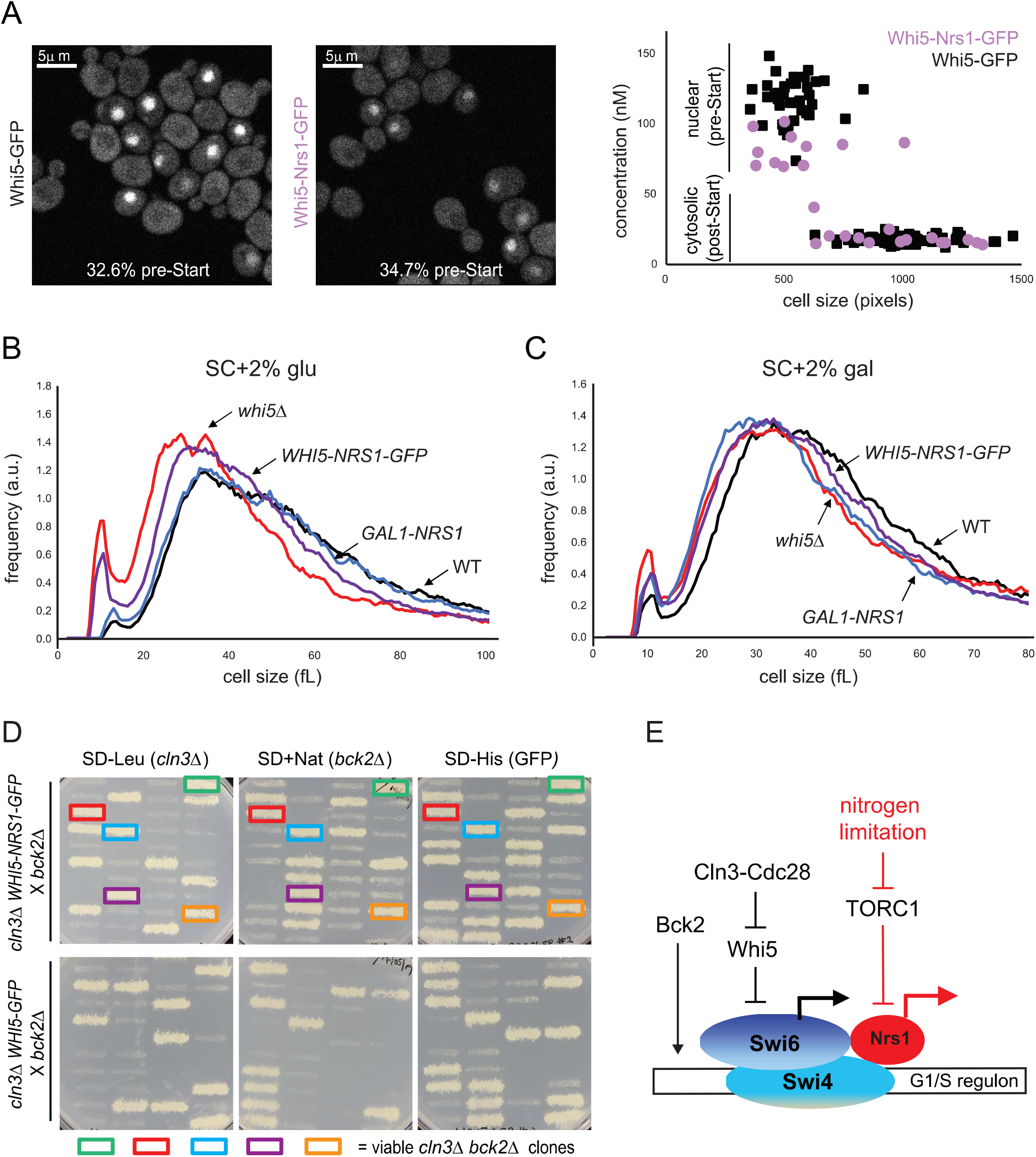
Tethered Nrs1 bypasses the Whi5-dependent lethal arrest of a *cln3Δ bck2Δ* strain. **(A)** sN&B microscopy images of *WHI5-GFP* and *WHI5-NRS1-GFP* cells grown in SC+ 2% glucose. Scale bars are 5 μm. The same intensity scale was used on both images. Fractions of pre-Start G1 cells were computed based on the assessment of nuclear GFP signal in *WHI5-GFP* cells (N=245) and *WHI5-NRS1-GFP* (N=95) cells. Absolute concentrations of Whi5-GFP and Whi5-Nrs1-GFP are shown in the plot. **(B)** Cell size distributions of wild type, GAL1-NRS1, whi5Δ and WHI5-NRS1-GFP strains in SC+ 2% glucose. **(C)** Cell size distributions of wild type, *GAL1-NRS1, whi5Δ* and *WHI5-NRS1-GFP* strains in SC+ 2% galactose. **(D)** Genotype of 10 tetrads from a *cln3Δ WHI5-NRS1-GFP* X *bck2Δ* cross (top) and a *cln3Δ WHI5-GFP* X *bck2Δ* cross (bottom). For each tetrad, spore clone growth was assessed on SD-LEU (indicates *cln3::LEU2*), SC+NAT (indicates *bck2::NAT*^*R*^), and SD-HIS (indicates *whi5::WHI5-NRS1-GFP-HIS3* or *Whi5::WHI5-GFP-HIS3*). Coloured boxes indicate viable *cln3Δ bck2Δ* spore clones, all of which also contained the *WHI5-NRS1-GFP* construct. (F) Simplified schematic for Nrs1-dependent activation of Start. Red lines indicate nitrogen-limited conditions. Not all components of the Start machinery are shown. See text for details.

As a reduction of Whi5 dosage in the Whi5-Nrs1-GFP fusion context could in principle explain the small size of this strain, we used sN&B to compare the nuclear concentrations of Whi5 in the wild type and the Nrs1-fusion contexts. Whi5 levels in wild type cells were 110-130 nM, consistent with those determined previously [31). The Whi5-Nrs1-GFP chimera was present at a slightly lower concentration, 80-l00nM (Figure 7A), close to endogenous Nrs1 nuclear levels in nitrogen-limited media (60-80nM, see Figure 2C). This slight decrease in Whi5 abundance was unlikely to explain the small size of cells expressing Whi5-Nrs1-GFP since cell size is not strongly sensitive to *WHI5* gene dosage [53]. We confirmed that hemizygous *WHl5/whi5Δ* diploid cells were only marginally smaller than wild type diploid cells (Figure S6E) and, moreover, *WHI5* overexpression only causes a 20-30% increase in mode size (Figure 3C), in agreement with previous results [53]. Consistently, our previously published Start model [31] also predicted that downregulating Whi5 levels from 120nM to 85nM should not affect the critical size at Start (Figure S6F).

Finally, we predicted that the Whi5-Nrs1 fusion should be sufficient to rescue the lethal Start arrest of a *cln3LJ bck2LJ* strain. We crossed a *cln3LJ* strain bearing the integrated *WHI5-Nrs1-GFP* allele to a *bck2LJ* strain and analyzed growth of dissected tetrads on selective media to identify spore genotypes (Figure 7D). We recovered many viable *cln3LJ bck2LJ WHI5-Nrs1-GFP* triple mutant spore clones but did not recover any viable *cln3LJ bck2LJ WHI5* double mutant clones (Figure 7D). As expected, no viable *cln3LJ bck2LJ* spore clones were obtained from a control cross of a *cln3LJ Whi5::WHI5-GFP* and a *bck2LJ* strain (Figure 7D). As a further control to ensure that the Nrs1 fusion did not merely inactivate Whi5 in a non-specific manner, we fused the above transcriptionally inactive C-terminal fragment of Nrs1 to Whi5 and showed that it did not reduce cell size (Figure S7A) nor rescue the *cln3LJ bck2’1* lethality (Figure S7B). These results supported a model in which Nrs1 bypasses Whi5 inhibition by directly conferring transactivation activity on the Whi5-inhibited SBF complex.

## Discussion

On the premise that additional genes may activate Start under suboptimal nutrient conditions, we screened for genes that can circumvent the Start arrest caused by loss of both *CLN3* and *BCK2* function. We discovered that *NRS1* overexpression efficiently bypasses the lethality of a *cln3LJ bck2LJ* strain and activates Start in wild type cells; that Nrs1 associates with SBF at G1/S promoter DNA and promotes SBF-dependent transcription; that *NRS1* overexpression can genetically suppress defects in SBF function; and that Nrs1 is itself a transcriptional activator. Our data suggest a model whereby nitrogen limitation or TORCl inhibition promote *NRS1* expression, likely via binding of the TORCl-regulated transcription factor Gln3 to the *NRS1* promoter [42, 43]. We hypothesize that Nrs1 binding to SBF result in direct recruitment of the transcriptional machinery such that Nrs1 bypasses Whi5-mediated inhibition of SBF (Figure 7E). In effect, endogenous Nrs1 acts as a nutrient-dependent parallel input into SBF. Although this input appears dispensable under most growth conditions, it is revealed in genetically sensitized context in laboratory strains and in wild yeast strains. We stress that the *cln3Δ bckΔ GAL1-WHI5* parental background used for the original high copy suppression screen might have biased towards the discovery of Start activators able to operate in presence of high Whi5 dosage.

This model of Nrs1 function (Figure 7E) allows re-interpretation of some previous *YLR053c/NRS1* genetic interactions uncovered in high-throughput studies [54]. Deletion of *NRS1* slightly aggravates the growth defect of a *ESS1* prolyl isomerase mutation which in turn exhibits negative genetic interactions with deletions of *SW/6* or *SW/4* [55]. Although the latter interactions suggested a possible role of Essl in Swi6/Whi5 nuclear import, we did not observe mis-localization of Swi4, Swi6 or Whi5 in either *GAL1-NRS1* or *nrs1Δ* strains. With respect to interactions with the transcriptional machinery, *NRS1* overexpression exacerbates the defect caused by deletion of *CTK1,* the catalytic subunit of RNA Pol II C-terminal domain (CTD) kinase I [56]. In contrast, *NRS1* deletion shows a positive genetic interaction with the RNA Pol II CTD-associated phosphatase, FCPl, which negatively regulates transcription [57]. *NRS1* overexpression also subtly increases chromosomal instability [58], which might result from premature Start activation [59]. These interactions with transcriptional regulators are consistent with our proposed model for Nrs1 function in G1/S transcription activation.

Nrs1 is a member of a newly described class of genetic elements variously called proto-genes, neo-ORFs, smORFS, small ORF-encoded peptides, or microproteins [37, 60–68]. Computational predictions and ribosomal footprinting first identified hundreds of short species-specific translated peptides from extragenic regions in yeast [37]. Subsequent approaches have yielded a plethora of microproteins encoded by smORFs in various species [60–63]. Documented functions for microproteins include phagocytosis [64], mitochondrial function [65], actin-based motility [66] and mitotic chromosome segregation [67]. De *nova* appearance of short proto-genes coupled with rapid evolution may form a dynamic reservoir for genetic innovation and diversification [37]. Interestingly, the rapid evolution of lncRNAs from junk transcripts has recently been reported [69], also consistent with the idea that regulatory functions can readily emerge de nova. Notably, fast evolving genes often have complex expression patterns compared to highly conserved genes and tend to be enriched for transcription-associated functions [70]. As illustrated by the example of *NRS1*, transcription activation may be a particularly facile route for proto-genes to quickly acquire important regulatory functions that optimize fitness under specific conditions.

## Methods

### Yeast strains construction and culture

All *S. cerevisiae* strains were isogenic with the S288C (BY4741) auxotrophic background (Supplemental Table 2). Standard molecular genetics methods were employed for genomic integration of C-terminal tagging cassettes [71]. Standard media were used for yeast growth: rich (XY: 2% peptone, 1% yeast extract, 0.01% adenine, 0.02% tryptophan); synthetic complete (SC: 0.17% YNB, 0.2% amino acids, 0.5% ammonium sulphate); or nitrogen-limited (0.17% YNB, 0.4% praline, supplemented with histidine, leucine, methionine and uracil to complement auxotrophies as needed). Prototrophic *S.boulardii* strains were grown in SC or nitrogen-limited (0.17% YNB, 0.4% praline) without amino acid supplements. Carbon sources were added to 2% w/v as indicated. Unless otherwise specified, cells were grown to saturation in SC+ 2% glucose and diluted 1 in 5000 into fresh medium for 16-18 h growth prior to experiments. These conditions ensured homeostatic growth in log-phase at the time of experiment at a culture density of 2-5 10^6^ cells/ml. For strains bearing constructs expressed under the inducible *GALl* promoter, pre-growth was in SC + 2% raffinose. Cell size distributions were acquired using a Beckman Z2 Coulter Counter. Growth curves were acquired at 30°C using a Tecan Sunrise shaker-reader.

### SGA screen

A *cln3Δ bck2Δ Whi5::GALl-WHI5* strain was mated to an array of 5280 *GALl-ORF* fusion strains [35]. Following haploid selection, strains were scored for growth on selective medium containing 2% galactose. Three replicates of this screen were performed. For the third replicate, expression of the *GAL4* transcription factor was increased in case it was limiting for expression of *WHI5,* the ORFs, or *GAL* genes required for growth in the presence of galactose at the sole carbon source. For this purpose, *GAL4* was placed under the control of the *ADHl* promoter in a *cln3::LEU2 bck2::NAT*^*R*^ *Whi5::Kan*^*R*^-*pGALl-WHI5 canld mfal::MFAlpr-spHIS5+ GAL4p::HphRpADH1-GAL4* query strain. Screen hits were validated by transformation of the *GALl-GST-ORF* construct directly into the *cln3Δ bck2Δ Whi5::GAL1-WHI5* query strain.

### Chromatin lmmunoprecipitation

Cells were grown in XY with 2% raffinose, induced with 2% galactose for 6 h to an OD_600_ ≤ 0.5 and fixed with 1% formaldehyde. Whole-cell extracts from 50 ml of culture were prepared by glass bead lysis, sonicated to shear chromatin DNA into fragments, and incubated with the appropriate antibody coupled to magnetic beads (Dynabeads PanMouse lgG). lmmunoprecipitated DNA was washed, de-crosslinked, purified, and analyzed by quantitative real-time PCR. Reactions with appropriate oligonucleotides were set-up with SYBR Green PCR Master Mix (Applied Biosystems) and carried out on an ABI 7500 Fast Real-Time PCR System. Enrichment at the *CLN2* or *PCLl* locus was determined after normalization against values obtained from input samples using *SYPl* as the reference gene. ^HA^Whi5 and Nrs1^13MYC^ ChlP experiments were performed in biological duplicate. The bar height on Figures SA and S4A represent the mean over duplicates, and the error bars represent the standard error on the mean.

### Immunoprecipitation and immunoblot analysis

Protein extracts were prepared in lysis buffer (10 mM HEPES-KOH pH 7.9, 50 mM KCI, 1.5 mM MgCl_2_, 1 mm EDTA, 0.5 mM DTT, 50 mM NaF, 50 mM sodium pyrophosphate, 1 mM Na_3_VO_4_ and Roche protease inhibitor cocktail) by glass bead lysis. Due to low expression levels in the 60-lS0nM range (Figure 2C and [31]), we immunoprecipitated proteins prior to immunoblot detection. lmmunoaffinity purifications were carried out at 4°C for 2 h with indicated antibodies, beads were washed 3 times with wash buffer (10 mM Tris-Cl pH7.9, 0.1% Triton X-100, 0.5 mM DTT, 0.2 mM EDTA, 10% glycerol, 150 mM NaCl), and resuspended in SDS sample buffer. Proteins were resolved by SDS-PAGE, transferred onto nitrocellulose membrane and immunoblotted with anti-HA, anti-Swi4 or anti-Swi6 (gift from Brenda Andrews), anti-GST or 9E10 monoclonal antibodies, followed by detection with HRP-conjugated secondary antibody [12, 72].

### In vitro binding assays

Recombinant GST-Nrs1 fusion protein was affinity purified with GSH-Sepharose 4B (Amersham Biosciences) in 50 mM HEPES-NaOH, pH 7.5, 150 mM NaCl, 5 mM EOTA, SmM NaF, 0.1% NP-40, 10% glycerol, supplemented with 1 mM PMSF. Recombinant SBF or MBF complex was purified with anti-FLAG resin (M2, Sigma) from insect cells co-infected with ^FLAG^Swi4-Swi6 or ^FLAG^Mbpl-Swi6 baculovirus constructs, in buffer supplemented with complete protease inhibitor cocktail (Roche), and then eluted with excess FLAG peptide [12]. Binding reactions were incubated at 4°C for 1 h with rotation. Washed samples were resolved on SOS-PAGE gel followed by immunoblotting with anti-FLAG (M2, Sigma) and anti-Swi6. SBF-GST-Nrs1 and SBF-HA-Whi5 complexes were pre-formed in solution and immobilized on glutathione or anti-HA resin, respectively, incubated with GST-Nrs1 or ^HA^Whi5, washed and analyzed as described above.

### Immunoprecipitation and mass spectrometry analysis

Cell pellets from untagged and Nrs1^13MYC^ strains from 100 ml of culture at 0D_600_ = 1 were lysed in standard lysis buffer (50 mM Tris-HCI pH8.0, 150 mM KCI, 100 mM NaF, 10% glycerol, 0.1% tween-20, 1 mM tungstate, 1 mM OTT, 10 mM AEBSF, 10 mM pepstatin A, 10 mM E-64) [SO] supplemented with protease inhibitors using a N_2_(I) freezer mill. Lysates (0.5 ml) were incubated for 1 h with 5 μL of anti-MYC antibody (Gentex) followed by 1 h incubation with an additional 50 μL of GammaBind plus Sepharose beads (GE Healthcare) to capture protein complexes. After multiple washes, samples were separated on BioRad precast gels and the entire gel lane was cut for each sample (CAPCA core facility, https://capca.iric.ca/; see Supplemental Methods). Gel samples were destained, alkylated and digested with trypsin for 8 h at 37°C and peptides extracted in 90% ACN. Peptides were separated on a home-made C18 column connected to Q-Exactive HF Biopharma with a 56 min gradient of 0% to 30% acetonitrile in 0.2% formic acid. Each full MS spectrum of extracted peptides was acquired at a resolution of 120,000, followed by acquisition of 15 tandem-MS (MS-MS) spectra on the most abundant multiply charged precursorions by collision-induced dissociation (HCD). Data were processed using PEAKS X (Bioinformatics Solutions, Waterloo, ON) and the UniProt yeast database. Variable selected posttranslational modifications were carbamidomethyl (C), oxidation (M), deamidation (NQ), acetyl (N-ter) and phosphorylation (STY). Control untagged and Nrs1^13MYC^ data were analyzed with Scaffold 4.8.9 at a false-discovery rate (FDR) of 1% for at least 2 peptides with a likelihood of at least 99%.

### RNA-seq analysis

Wild type and *GAL1-NRS1* strains were inoculated for overnight growth in SC+ 2% raffinose at 30°C, then diluted in SC+ raffinose to an OD600 of 0.07-0.1, incubated at 30°C until an OD600 of 0.15-0.2 was reached, at which point a sample was collected for RNA extraction prior to galactose induction. For *NRS1* induction, the cultures were either concentrated by centrifugation, resuspended in 20 ml SC+2% raffinose and added to 200 ml SC+2% galactose (replicate 2,3) or grown to an OD600 of 0.15-0.2 in 200 ml SC+ 2% raffinose followed direct addition of 2% galactose to the culture (replicate 1). Timepoint O was collected immediately, followed by timepoints up to 6 h. Total RNA was prepared as described in [73]. Briefly, cells were pelleted at 3500 r.p.m, disrupted using glass beads, phenol/chloroform extracted and the aqueous phase containing nucleic acids was ethanol precipitated. Contaminating genomic DNA was removed by DNase treatment (Qiagen). Total RNA was quantified using Qubit (Thermo Scientific) and RNA quality assessed with a 2100 Bioanalyzer (Agilent Technologies). Transcriptome libraries were generated using the Kapa RNA HyperPrep (Roche) using a Poly-A selection (Thermo Scientific). Sequencing was performed on the lllumina NextSeq 500 system with approximately 10 million single-end reads per sample. RNA-seq experiments were performed in biological triplicate.

Reads were aligned to all Ensembl yeast transcripts using Bowtie 2.2.5 [74] with default parameters. Read counts were tabulated for each gene, only considering alignments with an edit distance no greater than S. Genes with at least 500 reads in at least one sample were included in the analysis. Gene expression levels for each sample, expressed as log2 read counts, were normalized to the upper-quartile gene expression level for the same sample, as recommended in Bullard et al. [75], and represented as heatmaps Figure 5C. To control against sample to sample variability and the effects of medium change, gene enrichment scores were defined as follows. For each of the three *GAL1-NRS1* biological replicate samples, the expression after 6 h galactose induction was normalized separately to the expression in raffinose (prior to induction) and to the wild type sample from the same replicate after 6 h induction. From these, two log2(fold-change) scores were computed, and the score with the smallest modulus was kept as the final score for this replicat e. Score was set to zero if the signs of the two log2 fold-changes disagree d. Then, we averaged over the three replicates to get the final gene score, displayed in Supplemental Table 6. This procedure allowed to effectively control for the effects of the galactose media (normalizing by 6 h wild type samples), and for the inter-sample expression variability (through normalization to expression levels prior to induction and averaging over replicates). Hence, high scores for *GAL1-NRS1* (positive or negative) could be obtained only for genes up/ down-regulat ed both in galactose compared to raffinose, and compared to wild type cells in galactose. Wild type 6 h scores (Supplemental Table 6) were obtained exactly the same way, with the only difference that gene expression in wild type cells after 6 h in galactose were normalized separately to the wild type expression in raffinose and *GAL1-NRS1* after 6 h induction.

### High-content imaging

High-cont ent images were acquired on an OPERA high-throughput confocal microscope (PerkinElmer) equipped with a 60x water objective. Pixel resolution was 220 nm at 2*2 binning. 200 μL of culture were directly transferred to a Greiner Screenstar glass-bottom 96 well imaging plate and imaged within 30-45 min. GFP was excited with a 488 nm laser (800 ms exposure for endogenously tagged proteins, 320 ms for overexpressed proteins), and detected using a 520/35 nm band-pass filter. For *WHI5-GFP* and *WHI5-NRS1-GFP* strains, pre-Start G1 phase cells were identified by nuclear localization of the GFP signal, as computed using a custom image-analysis script written in MATLAB (MathWorks). The script masks individual cells using threshold-based detection of the autofluorescence background and determines the fraction of individual cells that display a pre-Start nuclear localization of the GFP signal.

### sN&B and R/CS fluorescence fluctuation microscopy

Live cells in log-phase were imaged on an Alba sN&B system (ISS, Champaign IL) comprised of an inverted confocal Nikon Eclipse microscope equipped with a lO0x water objective, a Fianium Whitelase continuous white laser with 488 nm emission filters and single photon APD detectors, as described previously [31]. In brief, 1.5 ml of cell culture was pelleted for 2 min at 3000 rpm, resuspended in ~100 μL of culture supernatant and 3 μL deposited on a pre-set 65 μL drop of SC+ 2% glucose+ 2% agar gel medium on a circular glass coverslip (#1, VWR) that was encircled by an adhesive silicon ring. After 4 min drying time, a ConcanavalinA-coated (Sigma, 2mg/mL) coverslip was gently pressed on top of the agarose to seal the pad against the adhesive silicon ring. Sealed pads were clamped in an AttoFluor chamber (Molecular Probes) and immediately imaged for no more than 1.5 h. Unless otherwise specified, the culture growth medium was re-used to make the imaging pads in order to prevent inadvertent nutrient up- or down-shifts. For this purpose, 1 ml of cell culture was pelleted at 15000 rpm for 1 min and 500 μL of the supernatant was mixed with 10 mg of agarose and warmed for 1-2 min at 98°C to melt the agar, followed by application to a coverslip. For sN&B experiments in the presence of rapamycin, growth medium containing 100 nM rapamycin was also used to prepare the pad. To mitigate possible degradation of rapamycin during the pad preparation, an additional 100 nM of rapamycin was added to the pre-warmed agar mix just before making the pad. The final rapamycin concentration was therefore 200 nM, i.e. about 200 ng/ml, similar to rapamycin treatment time-courses in Figure S1C and 4B, and the fixed-duration treatment in Figure 4A. Cells were pre-treated 1 h before imaging and imaging lasted about 1.5 h, yielding a treatment duration between 1 hand 2.5 h across the various fields of view (FOV). There were no significant differences in Nrs1-GFP signals across different FOVs.

sN&B imaging was performed using 20 raster scans of the same 30 μm-wide Fields of View (FOVs) of 256 pixels (pixel size 117 nm), using an excitation power of 1-2 μWat 488 nm wavelength and a 64 μs pixel dwell time. sN&B images shown in the figures are projections of the 20 raster scans. Protein concentrations were extracted from sN&B data using custom analysis software [31]. RICS imaging was performed in a similar fashion to sN&B imaging but different parameters were used to improve correlation curves (pixel size of 48.8 nm, 50 frames, 20 μs pixel dwell time) as described previously [50]. RICS vertical correlations for individual FOVs were fitted to single mode-free diffusion models using the SimFCS analysis software. FOVs of poor-quality fit were discarded from diffusion coefficients plots.

### Bioinformatics

The sequences of orthologs of *YLR053c* were obtained from the Fungal Orthogroups website. For S. *paradoxus* and S. *mikatae,* where no ORF was predicted at the expected locus, the *YLR053c* sequence was searched against whole genome assemblies using TBLASTN 2.2.27 to predict ort hologs. The synteny of the regions in *K. waltii* and S. *cerevisae* was confirmed based on Orthogroups orthology assignments of the upstream (47.17997 and *IES3*) and downstream (47.18006 and *OSW2*) genes and the fact that each of these assignments were supported by highly significant Blast e-values. Alignments were generated using ClustalO 1.2.0 and displayed using the MView online tool.

### Mathematical model of Start

Whisker and box plots for model predictions of the critical size at Start were obtained using the mathematical model and the data processing procedure described previously [31]. For each plot, 50 individual cells bearing variable Whi5 concentrations randomly picked in the 100-140 nM (for the *WHI5-GFP* strain) and 65-105nM (for the *WHI5-NRS1-GFP* strain) ranges were simulated, and plots generated in Excel using standard statistics of the distributions of critical size values for each strain.

## Data Availabiliy

The mass spectrometry data have been deposited to the ProteomeXchange Consortium via the PRIDE [76] partner repository with the dataset identifier PXD018681 and Doi 10.6019/PXD018681. The RNA-sequencing data have been deposited to the Gene Expression Omnibus database (GEO, NCBI).

## Acknowledgmens

We thank Marc Angeli for technical assistance with SGA screens, Brenda Andrews for providing reagents and strains and Elizabeth Bilsland for yeast fluorescent protein vectors. We also thank Jennifer Huber ofthe IRIC Genomics Platform for assistance with RNA-seq experiments. This work was supported by a grant from the Canadian Institutes of Health Research (FDN-167277) to M.T., a Genomics Technology Platform award from Genome Canada to M.T. and P.T., a Genomics Applications Partnership Program award from Genome Canada and Genome Quebec to P.T., an Institute for Data Valorisation (IVADO) postdoctoral fellowship to J.C.-H., and a Canada Research Chair in Systems and Synthetic Biology to M. T, and a Sigrid Juselius Foundation grant to S.T.

## Author contributions

Conceptualization: J.S., S.T., M.T.; Data Curation: S.T., J.C-H., E.B.; Formal Analysis: S.T., J.C-H., E.B.; Funding Acquisition: M.T., P.T., S.T.; Investigation and Validation: S.T., J.S., Y.T., R.P., G.G., X.T., S.M., D.B.; Methodology: S.T., J.S., R.P., X.T., S.M., C.A.R., M.T.; Project Administration and Supervision: S.T., M.T.; Resources: S.T., J.S., Y.T., R.P., G.G., X.T., S.M., D.B.; Software: S.T., J.C-H.; Visualization: S.T., J.S., Y.T., R.P., G.G., E. B.; Writing - Original Draft Preparation: S.T., J.S., D.B., E.B.; Writing-Review & Editing: S.T., Y.T., J.C-H., C.A.R., R.P., P.T., M.T.

