## Supplemental information for "A nitrogen source-regulated microprotein confers an alternative mechanism of G1/S transcriptional activation in budding yeast"

**for**

### Supplemental Methods

#### *Immunoprecipitation and mass spectrometry analysis*

Untagged and *NRS1*$^{13MYC}$ cell pellets from 100 mL of culture at $OD_{600}$ = 1 were lysed in lysis buffer supplemented with protease inhibitors using a freezer mill. Half of the final lysate (0.5 mL) was incubated for 1h with 50 µL of GammaBind plus Sepharose beads (GE Healthcare) and 5 µL of anti-MYC antibody (Gentex) to capture protein complexes. The other half was incubated with beads only and served as a negative control. Samples were washed four times and beads were resuspended in 50 µL of 2X lysis buffer. Samples were separated on BioRad precast gels and the entire gel line for each sample was prepared for mass spectrometry at the CAPCA core facility (IRIC, https://capca.iric.ca/). Gels were destained in 50% MeOH (Sigma-Aldrich), shrunk in 50% acetonitrile (ACN), reconstituted in 50 mM ammonium bicarbonate with 10 mM TCEP [Tris(2-carboxyethyl)phosphine hydrochloride; Thermo Fisher Scientific] and vortexed for 1 h at 37°C. Chloroacetamide (Sigma-Aldrich) was added for alkylation to a final concentration of 55 mM. Samples were vortexed for another hour at 37°C. Trypsin (1 µg) was added and digestion performed for 8 h at 37°C. Peptides were extracted with 90% ACN, dried down, solubilized in 5% ACN-0.2% formic acid (FA) and loaded on a C4 guard column (Optimize Technologies) connected directly to the switching valve. Separation was on a home-made reversed-phase column (150-µm i.d. by 180 mm) with a 56 min gradient from 10 to 30% ACN-0.2% FA at 600-nL/min flow rate on an Easy-nLC 1200 connected to an Q-Exactive HF Biopharma (Thermo Fisher Scientific, San Jose, CA). Each full MS spectrum acquired at a resolution of 120,000 was followed by 15 tandem-MS (MS-MS) spectra on the most abundant multiply charged precursor ions. Tandem-MS experiments were performed using collision-induced dissociation (HCD) at a collision energy of 27%. Data were processed using PEAKS X (Bioinformatics Solutions, Waterloo, ON) and the Uniprot yeast database. Mass tolerances on precursor and fragment ions were 10 ppm and 0.01 Da, respectively. Variable selected post-translational modifications were carbamidomethyl (C), oxidation (M), deamidation (NQ), acetyl (N-ter) and phosphorylation (STY). The data were visualized with Scaffold 4.8.9 (protein threshold, 99%, with at least 2 peptides identified and a false-discovery rate [FDR] of 1% for peptides). Hit proteins specific to Nrs1$^{13MYC}$ immunoprecipitates were obtained by subtracting proteins identified in the control sample and then filtering the candidate hits against the CRAPome database [1].

#### *Competitive growth assays*

Wild type (BY4741) and *nrs1::KanMX6* strains were transformed with 2 µm plasmids expressing Venus and mCherry from the *TDH3* promoter and bearing a *URA3* selection marker [2]. Single colonies of Venus and mCherry strains were inoculated in SC-URA + 2% glucose for 24h growth to saturation. Cell densities in each saturated culture were measured using a Beckman Z2 Coulter counter and an equal number of green and red cells of the two different strains (about 200µL of saturated culture per strain) were diluted in 5 mL of water. In parallel, an equal number of each individual strain was diluted in 5 mL of water as a fluorescence intensity control. 50 µL of each diluted culture was inoculated in triplicate into 5 mL SC-URA + 2%glucose or YNB + Leu + His + Met +0.4% Pro + 2% glucose, as indicated. Cultures were transferred to a 30°C rotary incubator and grown for 72h. Venus and mCherry fluorescence was measured for each culture with a Tecan M1000 plate reader (excitation 515nm and 587nm, respectively; emission 528nm and 610nm, respectively; 5 nm bandpass filters). The gain of the instrument was optimized using the single colour cultures to ensure signal linearity within the range of measurement. As a first approximation, the measured fluorescence signals in the red and green channels yield, respectively: $F_{red} = N_{red} * \varepsilon_{Cherry}$ , $F_{green} = N_{green} * \varepsilon_{Venus}$ , where $N_{red}$ and $N_{green}$ are the total number of red and green cells within the culture, and $\varepsilon_{Cherry}$ and $\varepsilon_{Venus}$ the amount of fluorescence emitted per cell in each channel. These equations can be rewritten in terms of the total number of cells in the culture, $N_0$, and the fractions $f_{red}$ and $f_{green}$ of red and green cells:

$$F_{red} = N_0 * f_{red} * \varepsilon_{Cherry} = N_0 * (1 - f_{green}) * \varepsilon_{Cherry}$$

$$F_{green} = N_0 * (1 - f_{red}) * \varepsilon_{Venus} = N_0 * f_{green} * \varepsilon_{Venus}$$

Hence, given a measurement of the fluorescence signals and the total number of cells in a mix, it is sufficient to know $\varepsilon_{Cherry}$ and $\varepsilon_{Venus}$ in all strain backgrounds. Coefficients were calculated in each strain background separately, using the single colour control cultures, where $f_{red}$ and $f_{green}$ = 0 or 1, and for which the residual fluorescence in the other channel (e.g., red channel for a Venus-coloured strain) was negligible (~0.1-0.5% at most). In principle, it is possible that a fluorescent protein *per se* may cause a change in cell fitness. To rule out this possibility, we also cross-analyzed complementary mixes, e.g., strain 1 tagged with mCherry and strain 2 tagged with Venus vs strain 1 tagged with Venus and strain 2 tagged with mCherry. For such mixes, if we denote $f_1$ the fraction of strain 1, the independence of $f_1$ with respect to colouring yields $F_{red,1} = N_{0,1} * f_1 * \varepsilon_{Cherry}$ and $F_{red,2} = N_{0,2} * (1 - f_1) * \varepsilon_{Cherry}$ from which we get:

$$f_1 = \frac{F_{red,1}/N_{0,1}}{F_{red,1}/N_{0,1} + F_{red,2}/N_{0,2}}$$

or, equivalently

$$f_1 = \frac{F_{green,2}/N_{0,2}}{F_{green,1}/N_{0,1} + F_{green,2}/N_{0,2}}$$

The data presented in Supplementary Figure S2E was derived using this analysis method, where each F/N ratio was averaged over 3 replicate cultures in each experiment. This method provided two measurements of the same fraction, based on analysis of the red and green fluorescence separately, and independently of renormalization to fluorescent signal in control cultures. With good accuracy (~2-3%), fractions derived using the green and red channels were identical. These values were also in good agreement with fractions derived using $\varepsilon_{Cherry}$ and $\varepsilon_{Venus}$ coefficients calculated in each strain background separately, using the single colour control cultures.


### Supplemental Figure Legends

**Supplemental Figure S1.** Additional characterization of Nrs1. **(A)** Full Ylr053c/Nrs1 protein sequence is conserved across the Saccharomyces sensu stricto group of species. Sequence alignment showing the Ylr053c/Nrs1 protein sequence in *S.cerevisiae* (top), aligned with sequences of orthologs in *S.bayanus* (c672-g32.1), *S.castellii* (656.13d), *S.paradoxus* and *S.mikatae* from top to bottom. Ylr053c/Nrs1 orthologs were not predicted in *S.paradoxus* or *S.mikatae* because of sharp length cutoffs in ORF prediction algorithms (the ORFs would span only 108 and 75 residues in *S.paradoxus* and *S.mikatae*, respectively). The lack of an obvious TATA box could also explain why no protein was predicted in *S.paradoxus*. Neighboring upstream and downstream genes both show high similarity to *YLR053c* neighbors in *S.cerevisiae*. The *YLR053c/NRS1* sequence also aligns in *S. kudriavzevii* (not shown). **(B)** Kinetics of Nrs1 expression upon nitrogen starvation. **s**N&B images of untagged wild type and *NRS1-GFP* log-phase cells (OD=0.4-0.7), following 22 h growth in YNB Pro medium from 1/5000 dilution (left) and 7 h growth from 1/100 dilution (right) from saturated pre-cultures. Arrows indicate representative Nrs1 signal beyond autofluorescence at 22h. **(C)** Prolonged exposure to rapamycin results in accumulation of a faster migrating form of Nrs1. Rapamycin was added to log-phase cultures of *NRS1*$^{13MYC}$ cells, aliquots were removed at indicated time intervals and immunoprecipitates were analyzed by anti-MYC immunoblot. **(D)** Nuclear localization of Nrs1 upon rapamycin treatment is not a consequence of the particular GFPmut3 fluorophore. Confocal microscopy image of *NRS1*$^{WT\text{-}GFP}$ cells grown in SC + 2% glucose medium, either untreated or treated with 200 ng/mL rapamycin for 2 h. **(E)** **s**N&B images of untagged wild type cells and *NRS1-GFP* cells grown to log-phase in SC + 2% glucose and plated on SC + 2% glucose agar pads containing either 0.5 M NaCl or 1 mM $H_2O_2$ and imaged over a 2 h time course. Images were acquired after ~1 h treatment, but Nrs1 expression was not observed at any time point for any of the treatments. **(F)** Nrs1 is not induced by DNA damage. *NRS1*$^{13MYC}$ cells grown in rich medium were exposed 0.1% MMS for 1 h prior to immunoprecipitation and *Nrs1*$^{13MYC}$ was detected with anti-MYC 9E10 antibody. Nrs1 expression from cells exposed to 200 ng/mL rapamycin and processed in parallel served as a positive control.

**Supplemental Figure S2.** Additional characterization of *nrs1Δ* strains. **(A)** Deletion of *NRS1* does not affect growth. Optical density (vertical axis) of wild type (black) and *nrs1Δ* (blue) strains grown in SC + 2% glucose (solid lines) or nitrogen-limited (YNB pro glu, dashed lines) medium as a function of time (horizontal axis). **(B-D)** Deletion of *NRS1* does not affect cell size. Cell size distributions of wild type (black) and *nrs1Δ* (blue) strains grown in SC + 2% glucose (**B**, solid lines), nitrogen-limited (**B**, YNB pro glu, dashed lines), SC + 2% galactose (**C**, solid lines), SC + 2% raffinose (**C**, dotted lines), YNB + 0.4% proline + 2% galactose (**C**, YNB pro gal, dashed lines), SC + 2% glucose (**D**, solid lines), SC + 4% glucose (**D**, dotted lines) and SC + 0.1% glucose (**D**, dashed lines). **(E)** Deletion of *NRS1* does not affect competitive fitness in SC + 2% glucose and nitrogen limited (YNB pro glu) medium during growth to stationary phase. Bar charts representation of the composition of 2 mixes of competing strains (Mix1:

WT transformed with mCherry plasmid (red) and *nrs1Δ* transformed with Venus plasmid (green); Mix2: WT with Venus plasmid (green), and *nrs1Δ* with mCherry plasmid (red)) as a function of time from inoculation. The percentage of each strain within the mixes shown is derived from 3 replicate cultures from the same original mixes (see Supplemental Methods). Error bars show the standard error on the mean. **(F)** *NRS1* is required for optimal growth in a strain that lacks MBF. Growth curves for wild type (BY4741), *nrs1Δ*, *mbp1Δ* and *mbp1Δ nrs1Δ* strains grown in nitrogen-limited YNB+Pro medium and nitrogen-rich SC medium. Curves represent the average of 3 different clones for each mutant strain. **(G)** *NRS1* is required for optimal growth in a wild yeast strain. Growth curves for wild type and *nrs1Δ S. boulardii* strains were grown in either nitrogen-poor YNB+Pro+0.1% glucose minimal medium (not supplemented with any other amino acids) or nitrogen-rich SC+0.1 % glucose medium. Three different isolates for the *nrs1Δ S. boulardii* strain were analyzed. Growth curves show are the average of at least 12 colonies for each strain.

**Supplemental Figure S3.** Example Swi4 and Swi6 peptide spectra detected in Nrs1 immunoprecipitates. The first of the five peptides identified for Swi4 (ITSPSSYNKTPR) and Swi6 (SGLRPVDFGAGTSK) are shown on top and bottom, respectively. Data were processed with Scaffold software.

**Supplemental Figure S4.** Lack of effect of Nrs1 on Whi5. **(A)** *NRS1* overexpression does not inhibit Whi5 association with G1/S promoter DNA. Wild-type or *WHI5*$^{HA}$ strains carrying empty vector or a *GAL1-NRS1* plasmid were grown in SC + 2% raffinose medium and induced with 2% galactose for 6 h prior to crosslinking. Anti-HA chromatin immunoprecipitations were assessed for the presence of *CLN2* and *PCL1* promoter DNA by quantitative RT-PCR. Bars indicate the mean fold-enrichment across two replicates, error bars show the standard error on the mean. **(B)** *NRS1* overexpression does not affect Whi5 protein levels. Whi5-GFP absolute concentration in single wild type (blue dots) and *GAL1-NRS1* (orange dots) cells first grown in SC + 2% raffinose then induced with 2% galactose for 6 h prior to sN&B microscopy. Nuclear Whi5-GFP levels in pre-Start cells and cell-averaged levels in post-Start cells where Whi5 has been exported from the nucleus are shown. **(C)** *NRS1* overexpression does not inhibit Whi5 association with SBF. The indicated Whi5$^{HA}$ immunoprecipitates from strains induced with galactose for 6 h were probed for Whi5$^{HA}$, Swi4, or Swi6 by immunoblot. **(D)** Nrs1 does not compete with Whi5 for binding to SBF *in vitro*. The indicated amounts of recombinant $^{HA}$Whi5 or $^{GST}$Nrs1 was titrated into preformed $^{FLAG}$Swi4-Swi6-$^{GST}$Nrs1 or $^{FLAG}$Swi4-Swi6-$^{HA}$Whi5 complexes immobilized on anti-FLAG resin, respectively. Bound proteins were resolved by SDS-PAGE then immunoblotted (top) or stained with Coomassie Brilliant Blue (bottom). Note that added soluble $^{GST}$Nrs1 or $^{HA}$Whi5 saturated the respective SBF-$^{HA}$Whi5 and SBF-$^{GST}$Nrs1 complexes at the lowest input concentrations.

**Supplemental Figure S5.** Additional data relevant to Nrs1-mediated transactivation and *GAL1-NRS1* genetic interactions. **(A)** Control growth curves for trans-activation assays. Reporter strains transformed with plasmids expressing either *GAL4*$^{DBD}$ alone, *GAL4*$^{DBD}$*-NRS1*, *GAL4*$^{DBD}$*-UBE2G2* or *GAL4*$^{DBD}$*-NRS1*$^{Cter}$ were grown in SD-Trp medium at 30$^{o}$C. **(B)** *NRS1* overexpression does not rescue a *cln1Δcln2Δcln3Δ* G1 phase arrest. Left: Cultures of *cln1Δcln2Δcln3Δ MET-CLN2* and *cln1Δcln2Δcln3Δ MET-CLN2* + *<pGAL1-NRS1>* strains grown to log-phase in SC-Met+2% raffinose, then reinoculated in either SC-Met+2% raffinose, SC-Met+2% galactose, SC+Met+2% raffinose or SC+Met+2% galactose for the indicated periods of time before determination of cell size distributions on a Beckman Z2 Coulter counter. Right: bar charts showing the average number of cell divisions for *cln1Δcln2Δcln3Δ MET25-CLN2* and *cln1Δcln2Δcln3Δ MET25-CLN2* + *<pGAL1-NRS1>* strains during the 18 h interval between the 6 h and 24 h timepoints in SC+Met+2%galactose. Bar heights represent the average of 4 different clones (N=4); error bars represent the standard deviation. **(C)** Room temperature growth controls for genetic interactions of *NRS1* with *SWI4* and *MBP1*. Serial 5-fold dilutions of wild type *NRS1* and *nrs1*::*GAL1-NRS1* strains in wild type (rows 1, 2, 10), *swi4-ts* (row 3), *mbp1Δ* (row 4) and *mbp1Δswi4-ts* (rows 5-9) backgrounds were spotted onto SC + 2% glucose, SC + 2% raffinose and SC + 2% galactose medium and grown for 5 days at 23°C. C1-4 are four clones of *mbp1Δ swi4-ts GAL1-NRS1*. **(D)** Images of the same serial 5-fold dilutions of *NRS1* and *GAL1-NRS1* strains in wild type, *swi4-ts*, *mbp1Δ* and *mbp1Δ swi4-ts* backgrounds as in Figure 6B, spotted onto SC + 2% glucose, SC + 2% raffinose and SC + 2% galactose, but grown for an additional 2 days (i.e., 7 days total growth time at 30°C). C1-C4 are four clones of *mbp1Δ swi4-ts GAL1-NRS1*.

**Supplemental Figure S6.** Additional characterization and controls for Nrs1 and Whi5 fusion proteins. **(A)** The Whi5-Nrs1-GFP chimeric protein is produced *in vivo* and migrates at an expected size. *WHI5-GFP*, *NRS1-GFP* and *WHI5-NRS1-GFP* strains were grown in nitrogen-limited (YNB+pro) medium and extracts immunobloted with anti-GFP antibody. **(B)** A C-terminal fusion of Nrs1 to Whi5 does not affect cell cycle distribution. High-content images of *WHI5-GFP* and *WHI5-NRS1-GFP* cells grown in SC

+ 2% glucose were acquired on an OPERA high-throughput confocal microscope (PerkinElmer) equipped with a 60x water objective. The same intensity scale was used for both panels. Scale bar is 10 μm. The fraction of pre-Start (G1) cells was obtained using a custom MATLAB script (see Methods). **(C)** Fusion of a GFP tag at the Whi5 C-terminus does not affect cell size. Cell size distributions of untagged wild type and *WHI5-GFP* cells grown in SC + 2% glucose were determined on a Beckman Z2 Coulter counter. **(D)** Growth curves of wild type, *whi5Δ*, and *WHI5-NRS1-GFP* strains in SC + 2% glucose medium at 30°C. **(E)** *WHI5* dosage has only minor effects on cell size. Cell size distributions of wild type and *WHI5/whi5* heterozygous diploid strains grown in SC + 2% glucose. Dotted, dashed and solid blue lines represent three different *WHI5/whi5* clones. **(F)** Predicted effects of *WHI5* dosage on cell size in a mathematical model of Start. Box and whisker plots show distribution of critical cell sizes predicted by the Start model published in [3] for simulated average Whi5 concentrations of 120 nM (corresponding to *WHI5-GFP* cells, left boxplot) and 85 nM (corresponding to *WHI5-NRS1-GFP* cells, right boxplot).

**Supplemental Figure S7.** Nrs1 function at G1/S requires the poorly conserved N-terminal region. **(A)** Cell size distributions of wild type and *WHI5-NRS1$^{Cter}$-GFP* strains grown in SC+2% glucose determined on a Beckman Z2 Coulter counter. **(B)** Genotype of 10 tetrads from a *cln3Δ whi5::WHI5-NRS1$^{Cter}$-GFP* X *bck2Δ* cross. For each tetrad, spore clone growth was assessed on SD-Leu (indicates *cln3::LEU2*), SC+NAT (indicates *bck2::NAT$^{R}$*), and SD-HIS (indicates *whi5::WHI5-NRS1$^{Cter}$-GFP-HIS3*). Blue boxes indicate viable *cln3Δ* or *bck2Δ* spore clones. No viable *cln3Δ bck2Δ* double mutant clones were recovered.

### List of Supplemental Tables

**Supplemental Table 1.** Candidate dosage suppressors of *cln3Δ bck2Δ* lethality.

**Supplemental Table 2.** Yeast strains and plasmids used in this study**.**

**Supplemental Table 3.** Proteins detected specifically in Nrs1 immunoprecipitates by mass spectrometry.

**Supplemental Table 4.** List of peptides identified in Nrs1 and control immunoprecipitates by mass spectrometry.

**Supplemental Table 5.** List of proteins identified in Nrs1 and control immunoprecipitates by mass spectrometry.

**Supplemental Table 6.** Calculated gene scores for RNA-seq experiments performed with wild type and *GAL1-NRS1* strains. For upregulated (score > 0) and strongly upregulated (score > 0.5) genes, Swi4/Mbp1-binding scores were imported from Ferrezuelo et al. [10], and SBF/MBF targets were identified and counted. This data was used to perform the hypergeometric enrichment tests.

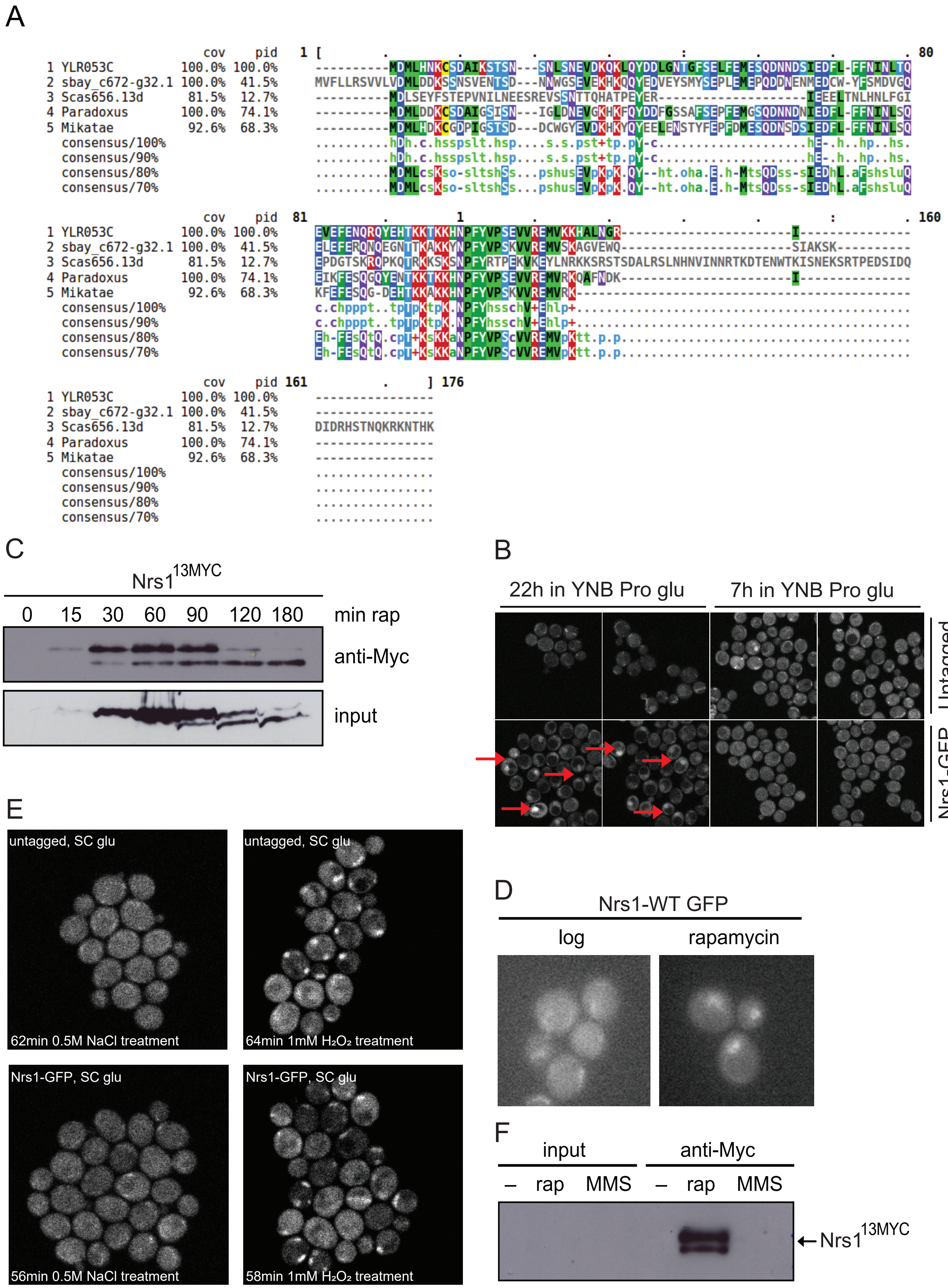


**Tollis *et al.* Supp. Figure S1**

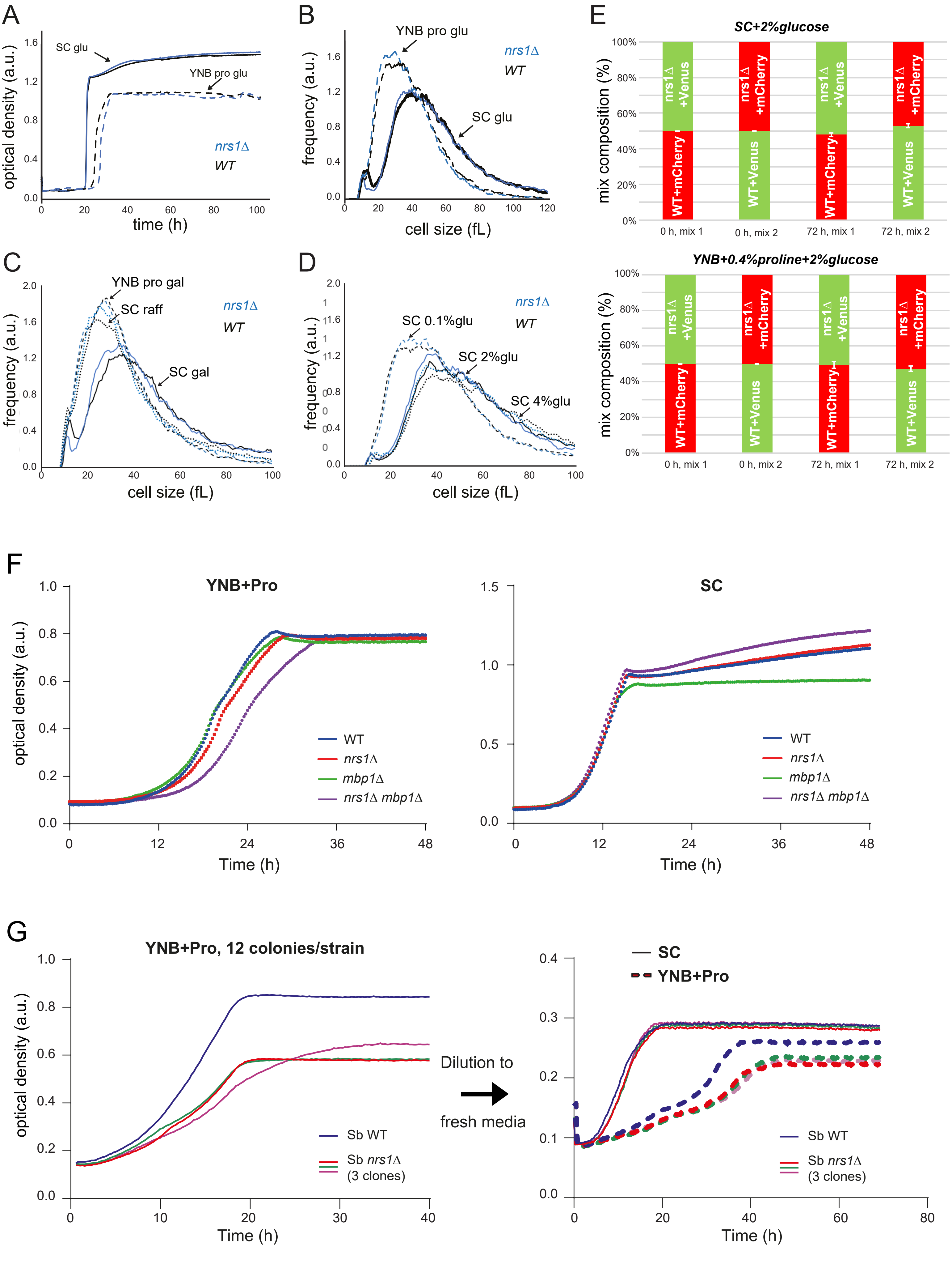


**Tollis *et al.* Supp. Figure S2**

A

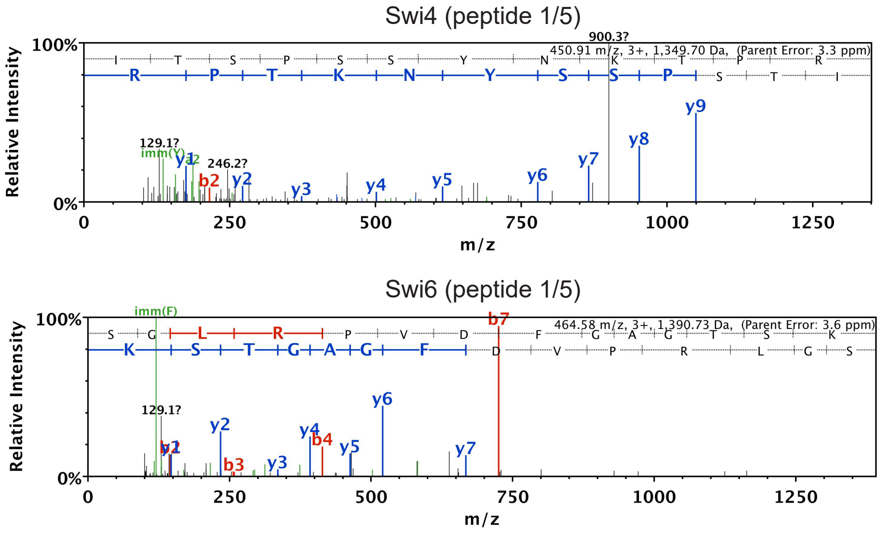


**Tollis *et al.* Supp. Figure S3**

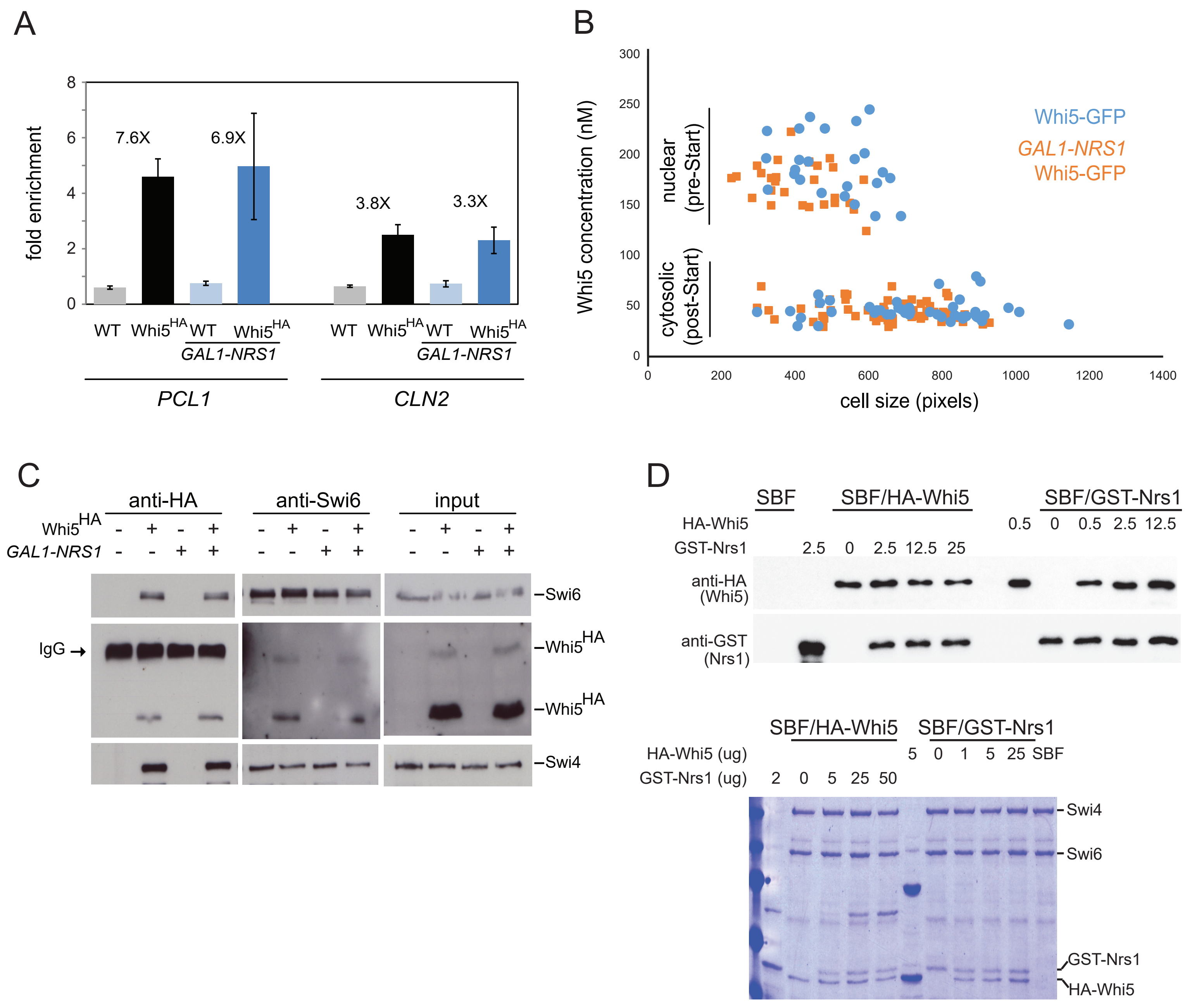


**Tollis *et al.* Supp. Figure S4**

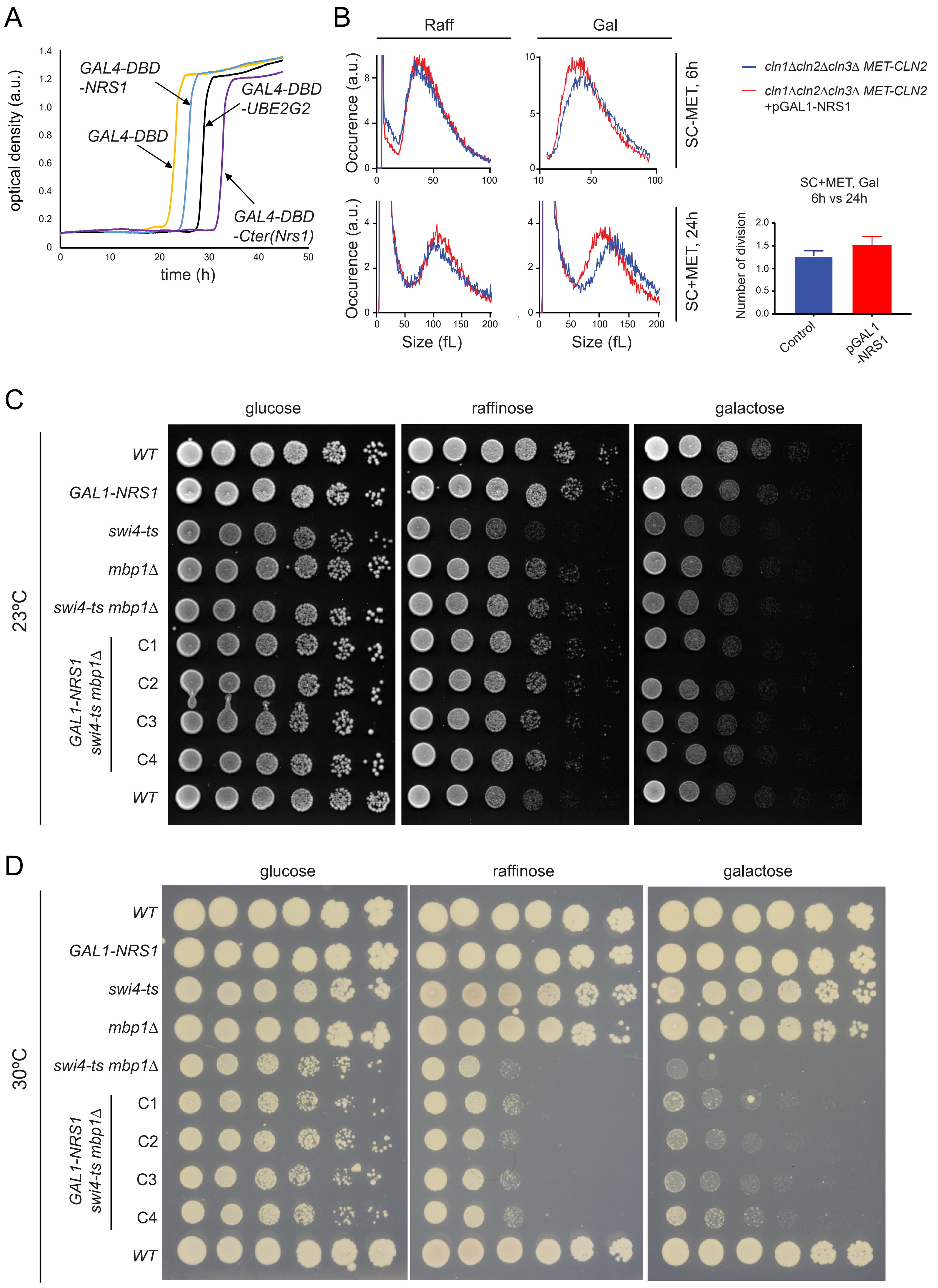


**Tollis *et al.* Supp. Figure S5**

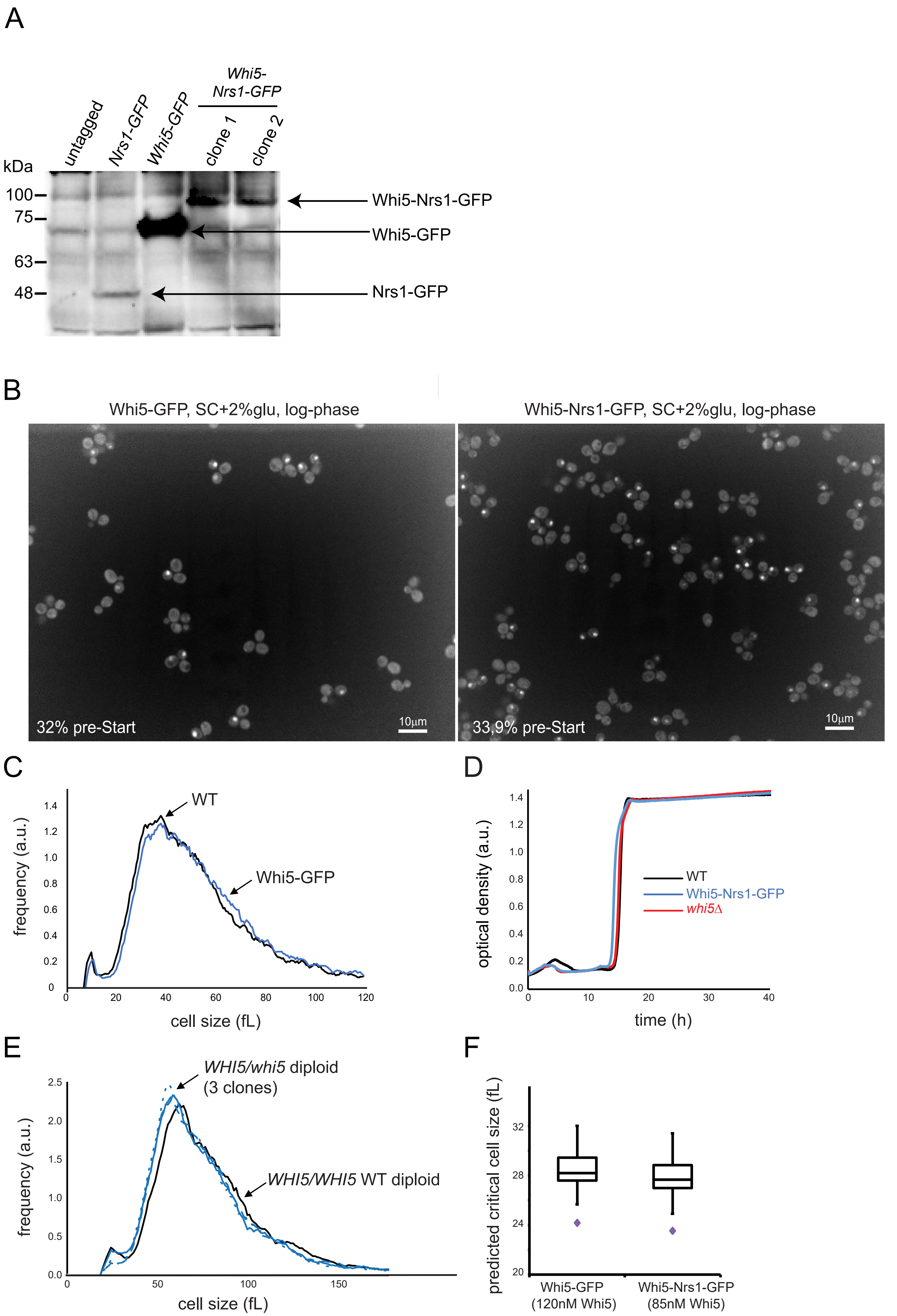


**Tollis *et al.* Supp. Figure S6**

A

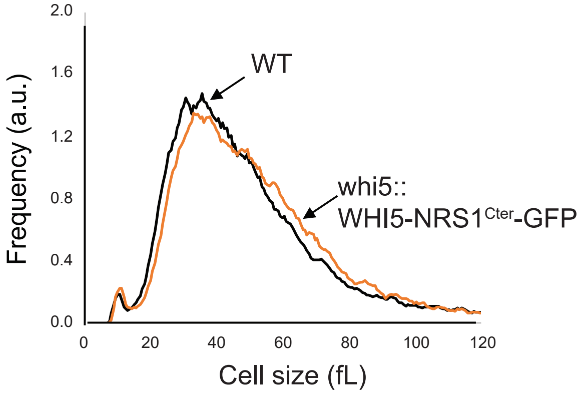


B

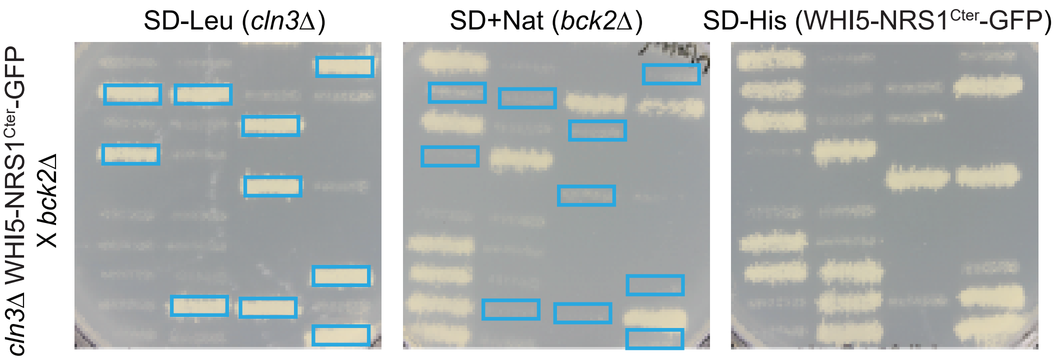


**Tollis *et al.* Supp. Figure S7**
